# SWI/SNF ATPase Brahma and Notch signaling collaborate with CBP/p300 to regulate neural stem cell apoptosis in *Drosophila* larval central nervous system

**DOI:** 10.1101/2025.06.16.659888

**Authors:** Punam Bala, Viswadica Prakki, Rohit Joshi

**Affiliations:** Laboratory of Neuroscience and Cell Biology, Centre for DNA Fingerprinting and Diagnostics (CDFD), BRIC-CDFD, Inner Ring Road, Uppal, Hyderabad-500039. India; Graduate studies, Regional Centre for Biotechnology, Faridabad 121001

## Abstract

SWI-SNF ATPase Brahma and Notch signalling are known to interact during development, but how this interaction is executed at the molecular level is not fully understood. We have investigated the molecular mechanism of Brm-Notch interaction in the context of Hox-dependent neural stem cell (NSC) apoptosis in the developing Central Nervous System (CNS) of *Drosophila*. Our results suggest a multi-tier regulation of NSC apoptosis by Brahma, first by regulating the expression of Drosophila CBP/p300 (Nejire) and the molecular triggers of cell death (Hox, bHLH factor Grainyhead, and Notch signalling pathway). The second mode of regulation is by direct binding of Brahma to the apoptotic enhancer and its collaboration with Notch signalling pathway to regulate the *RHG* family of apoptotic genes, *grim and reaper*. Our data support a model where, upon activation of Notch signaling, Brahma and CSL-Su(H)/Mastermind complex recruit CBP/p300 onto the apoptotic enhancer. This increases the H3K27ac marks on the nucleosomes to open up the chromatin and facilitate apoptotic gene transcription in Abd-B and Grh dependent manner.

## Introduction

Precision of the neural circuitry in the developing central nervous system (CNS) of bilaterian organisms critically relies on generating the right cellular diversity [1]. The generation of this diversity and the maintenance of tissue homeostasis in a region-specific manner require a fine balance of proliferation, differentiation and apoptosis of neural stem cells, neurons and glia [2, 3]. This is directly and indirectly orchestrated in a segment-specific manner by the Hox family of transcription factors (**TFs**) [4–23]. Programmed cell death (**PCD**-apoptosis) plays a particularly important role in the patterning of the CNS, with neuronal and glial apoptosis being critical for the refinement of the neuronal circuitry [24–26] and neural stem cell apoptosis for bringing about gross patterning changes in the CNS [4, 6, 13, 14, 17, 20–23, 27–29].

During the developmental morphogenesis of the CNS, the apoptotic triggers range from loss of survival signals, transcriptional induction of pro-apoptotic genes, growth factor signalling, intrinsic transcription factors or a combination of two or more of these events. The transcriptional induction of pro-apoptotic genes is used as a trigger in both vertebrates [30, 31] and invertebrates [8, 10, 20–25]. Segment-specific Hox-dependent neural stem cell (NSCs) apoptosis observed in *Drosophila* CNS is an example of one such mode of cell death regulation in developing CNS, which is crucial for tissue homeostasis and developmental morphogenesis [4, 6, 10, 13, 14, 17, 21–23]. This mode of NSC (or Neuroblasts –**NBs**) apoptosis happens across the CNS [4, 6, 13, 14, 17, 20–23, 27–29], and has been shown to rely on the resident Hox protein and its cofactor Grainyhead (**Grh**) in larval stages [6, 20, 22, 23, 29]. In the case of abdominal (A3-A7) and terminal (A8-A10) segments of the larval CNS the cell death is initiated in mid third instar (**mL3**) stage (Fig. 1A), when Grh and the resident Hox factors (expressed in these regions, Abdominal-A and Abdominal-B, respectively) collaborate with Notch signalling pathway to transcriptionally activate the *RHG* (***reaper, hid and grim***) family of apoptotic genes [32, 33]. This activation happens through a 1 kb enhancer (***F3B3***), deletion of which results in block of NB apoptosis (Fig-1A)[20, 22, 23].

**Figure 1:**
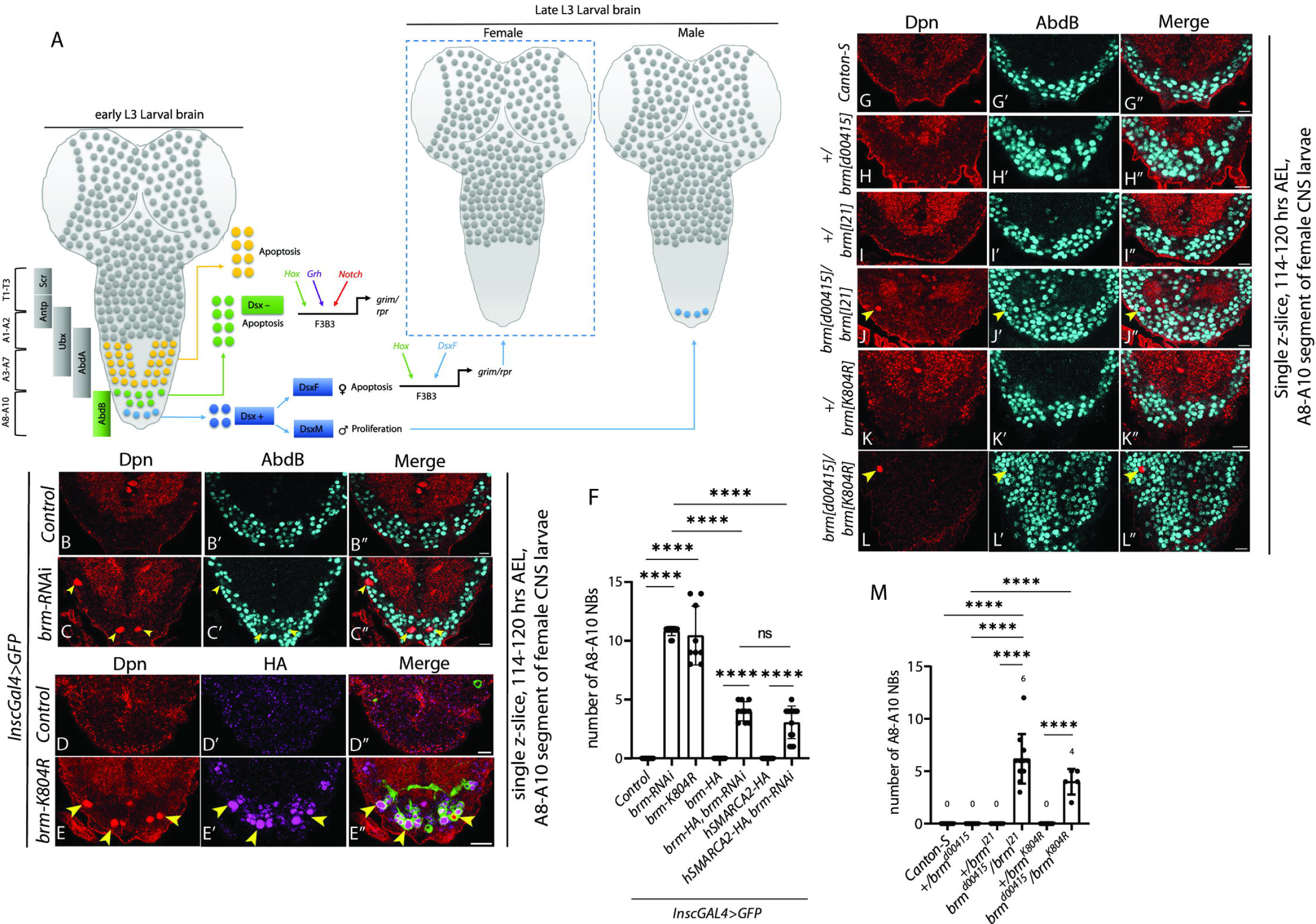
ATPase activity of Brm is required for A8-A10 NB apoptosis. **(A)** Schematic of larval CNS showing Hox-dependent NB apoptosis in abdominal (A3-A7) and terminal (A8-A10) segments. In Abd-B expressing terminal (A8-A10) segments, NBs are categorized based on the expression of *doublesex* (*dsx*). In female CNS, Dsx-positive NBs use Abd-B and Dsx^F^ (female-specific isoform) to undergo apoptosis, while in males, these NBs continue dividing till late pupal stages. Dsx-negative NBs (hereon referred to as A8-A10 NBs) use Hox (Abd-B), Grh and Notch signalling to transcriptionally activate RHG family of apoptotic genes (*grim* and *reaper*) through a 1 kb enhancer (*F3B3*) and intiate apoptosis in mid L3 stage. **(B-E)** Shows terminal segments of late L3 stage CNS with no surviving (A8-A10) NBs in case of controls (*inscGAL4>GFP*) (B, D) (0±0 NBs, *n*C:=C:8 VNCs, N□*=*□*2*) and ectopic NBs in *brm-RNAi* (10.83±0.13 NBs, *n*C:=C:12 VNCs, N = 3) and over-expression of HA-tagged Brm-K804R (10.44±0.88 NBs, *n*C:=C:9 VNCs, *N*C:=C:3). This establishes that *brm* knockdown blocks NB apoptosis, and ATPase activity of Brm is required for this apoptosis. **(F)** Graph showing the number of surviving A8-A10 NBs in the case of CNS of controls, *brm-RNAi,* and *brm-K804R*. Graph also shows the number of surviving NBs when the *brm-RNAi* phenotype was rescued using overexpression of Drosophila Brm (*HA-brm*; 4.00±0.29 NBs, *n*C:=C:10 VNCs, *N*C:=C:4), and the human ortholog of *brm (HA-hSMARCA2;* 3.07±0.49 NBs, *n*C:=C:14 VNCs, *N*C:=C:4) (Bars 5, and 7), establishing cross-phylum conservation of the Brm function. **(G-L)** Shows that no NBs survive in CNS of controls (*Canton S*) and heterozygotes for *brm* alleles (*brm^d00415^/+* and *brm^I21^/+*; *brm^K804R^/+;* 0±0 NBs, *n*C:=C:7 VNCs, *N*C:=C:2), but NBs apoptosis is blocked in hetero-allelic combinations of *brm* (*brm^d00415/I21^*; 6.18±2.35 NBs, *n*C:=C:11 VNCs, *N*C:=C:4; *brm^d00415^*^/*K804R*^; 4.00±1.22 NBs, *n*C:=5 VNCs, *N*C:=C:2). **(M)** Graph showing the number of surviving A8-A10 NBs in heterozygous controls and heteroallelic *brm* mutant combinations. Yellow arrowheads indicate the ectopic NBs. Scale bars are 10 µm. Graph shows mean±SD. For zero only control and non-zero test, significance is from Fisher’s test and for non-zero test and control, significance is from a Generalized Linear Model (GLM) with a binomial logistic regression performed using *‘R’.* “n” shows the number of VNC analysed, and “N” is number of times the experiment was repeated in the current and all subsequent figures.

Notch signalling pathway is an evolutionarily conserved, juxtacrine mode of signalling between adjacent cells [34]. The Notch receptor is localised on the membrane of the signal-receiving cell. Upon the activation of Notch by membrane-bound ligands like Delta/Serrate, the Notch intracellular domain (**NICD**) gets cleaved and goes inside the nucleus and binds to the executive transcription factor CSL (or Suppressor of Hairless (**Su(H)** *in Drosophila*) [34]. Following this event, co-repressor binding to Su(H) is replaced by co-activator Mastermind (***mam***), which recruits CBP/p300, a Histone AcetylTransferase (**HAT** encoded by *nejire-**nej*** in *Drosophila*) [34, 35], resulting in downstream target gene activation. Expectedly, Notch signalling activation has been shown to bring about CBP/p300-dependent increase in H3K56ac marks and associated chromatin accessibility changes on its target enhancers [35]. In *Drosophila* salivary glands, the enhancer accessibility preceding the signalling activation is shown to be facilitated by SWI-SNF ATPase Brahma (**Brm**), which Su(H) bind onto the enhancer DNA [36], though this interaction needs to be validated in a developmentally specialised tissue-specific context. Prior studies have also shown a connection between Brm and Notch signalling pathway, with Brm-BAP complex regulating Notch pathway components and its targets [36, 37], similarly, a dominant interaction of Delta and the ATPase dead version of Brm has also been reported [38]. Even though there is evidence of interaction between Brm and Notch signalling during development, the molecular details of how this interaction is effected are not fully understood.

In this work, we investigated Brm and Notch interaction in the context of Hox-dependent NB apoptosis in the larval CNS of *Drosophila*. We find that Brm controls NB apoptosis in terminal segments of the CNS by regulating the expression of Hox (Abdominal-B, **Abd-B**), Grh, and the Notch activity in the NBs. The regulation of the Notch signalling happens at two levels. First, Brm regulates the expression of members of the Notch signalling pathway (like the Notch receptor and its co-activator Mastermind). Second tier of regulation is at the level of the apoptotic enhancer, where Brm interacts with Notch signalling to activate the expression of apoptotic genes *grim* and *reaper*; which is corroborated by an in vitro physical interaction between Su(H) and Brm. The NB apoptosis also required the *Drosophila* ortholog of CBP/p300 (Nejire-Nej) at the apoptotic enhancer. Congruent with this, we found that Brm could successfully pull down Su(H), Mam and Nej from larval lysates.

Collectively, our results support a model where Brm regulates the enhancer accessibility and activation of apoptotic genes, indirectly by modulating the expression of Hox, Grh, CBP/p300 and the members of the Notch signalling pathway, and directly by collaborating with Su(H)/CBP signalling complex on the apoptotic enhancer.

## Results

### ATPase activity of Brahma is required for NB apoptosis in larval CNS

The NBs in abdominal (30 in A3-A7) and terminal (20 in A8-A10) segment of the larval ventral nerve cord (**VNC**) start dying in second instar stage and are completely eliminated from VNC by late L3 stage (**LL3**) [22, 23]. In the terminal segments, NBs are classified based on the expression of the *doublesex* gene (*dsx*) [22]. In female CNS, four Dsx-positive NBs undergo sex-specific apoptosis (which is independent of Grh and Notch) in mid L2 stage of development [21](Fig. 1A). Their counterparts in male CNS continue dividing till pupal stages. The remaining 16 Dsx-negative NBs start dying in mL3 stage [22] and are studied and analysed here.

Since SWI-SNF ATPase Brahma (*brm*) is known a regulator of Hox genes, we wanted to test if it played a role in Hox-dependent NB apoptosis. Continuing on this thinking, we tested Brm and members of P/BAP complex using RNA interference (RNAi) for candidate genes (Table-1) regulating NB apoptosis in abdominal and terminal segments of larval ventral nerve cord. We tested five RNAi lines targeted against different regions of the Brm cDNA and observed that three of these lines (detailed in methods and Table-1) showed a block of NB apoptosis in larval VNC. Owing to better consistency of the expression of the previously characterised *enhancer-lacZ* line (*F3B3-lacZ*), and the fact that the *brm* knockdown was unable to block the apoptosis of Dsx-positive NBs in the female CNS, we focused our experiments and analysis on Dsx-negative NBs found in the A8-A10 segments of the female larval CNS. These cells were identified by costaining with the NB marker Deadpan (Dpn) and the resident Hox factor Abd-B, and none of these NBs survive by the late L3 stage in female CNS [21, 22].

We observed that *brm*-knockdown (using NB expressing *inscGAL4*) could block the death of Dsx-negative NBs (Fig. 1B-1C and 1F). Since making a z-project for all the slices compromises the signal-to-noise ratio and obscures information, single confocal z-slices are shown as representative images (Fig. 1B-1C) in this and subsequent experiments, and the quantification of the data is shown in the graph (Fig. 1F). All three lines blocked apoptosis when the knockdown was induced from the embryonic stages. We also observed a block of NB apoptosis in the case of Moira and Snr1 (two RNAi lines each), which are components of the Brm-associated core complex (Fig. S1C-S1F). Thereafter, we did a rescue experiment to establish the specificity of the phenotype and conservation of the function of the *brm* gene across species. We could partially rescue the block of NB apoptosis by overexpression of the *Drosophila brm* gene, as well as with its human ortholog (hSMARCA2) (Fig 1F and Fig. S1U-S1V). To further confirm the phenotype, we tested a weak heteroallelic mutant combination for the hypomorphic alleles of *brm* gene (*brm^d00415/I21^*), which survived till late third instar larval stage (LL3) of development, and we could observe a block of NB apoptosis in this case as well (Fig. 1J and 1M). Next, we tested the role of the BAP and PBAP complex in NB apoptosis. Here, we observed that knockdown for Osa, which is a BAP complex specific subunit, blocked the NB apoptosis (Fig. S1J-S1K); while the knockdown for PBAP complex-specific subunits (like SAYP, BAP170, and Polybromo) did not (Fig. S1L-S1Q).

Thereafter, we tested the contribution of Brm’s chromatin remodelling activity in NB apoptosis. The remodelling activity of Brm relies on the ATPase domain, and mutating lysine-804 to an arginine (K804R) is known to abrogate it [39–41]. We observed that overexpression of the ATPase domain dead version of the Brm (Brm-K804R) also blocked NB apoptosis in A8-A10 segments (Fig. 1E and 1F), comparable to what was seen in the case of *brm-RNAi*. To conclusively establish the role of K804 in NB apoptosis, CRISPR-Cas9 was used to mutate lysine 804 to arginine in *brm* (*brm^K804R^*). A weak heteroallelic mutant combination of the CRISPR allele and hypomorphic allele (*brm^d00415/K804R^*) survived until the LL3 stage of development and also showed a block of NB apoptosis (Fig. 1L and 1M).

These results established that the Brm and members of BAP complex are important in executing Hox-dependent NB apoptosis and which relies on Brm’s

ATPase activity for this function. The rescue of the block of NB apoptosis by the human ortholog hSMARCA2 suggests a conservation of gene function across species.

### Brm regulates NB apoptosis by regulating Hox, Grh and Notch signalling pathways

Next, we checked the levels of Grh, Hox and Notch activity (which function as triggers for A8-A10 NB apoptosis) for control and in the case of *brm* knockdown.

Since these NBs asynchronously initiate apoptosis by the mid-L3 stage and are cleared from the VNC soon after, it is difficult to consistently monitor target gene (or *F3B3-lacZ*) expression in these cells in the late L2 or early L3 stage. To circumvent this problem, we used larvae in which NB apoptosis was blocked by expression of the cell death blocker p35 as controls, and compared target gene (or *F3B3-lacZ*) expression in these cells with that in age-matched test larvae (brm knockdown) at the late L3 stage. The Notch activity was monitored using E(spl)mγ-GFP, which shows a strong expression and is known to be responsive to Notch signalling in NBs [42]. We observed that *brm*-knockdown resulted in a significant reduction in the levels of Abd-B (Fig. 2B’), Grh (Fig. 2E’) and a complete loss of E(spl)mγ-GFP (Fig. 2H’) in the NBs in A8-A10 segments.

**Figure 2:**
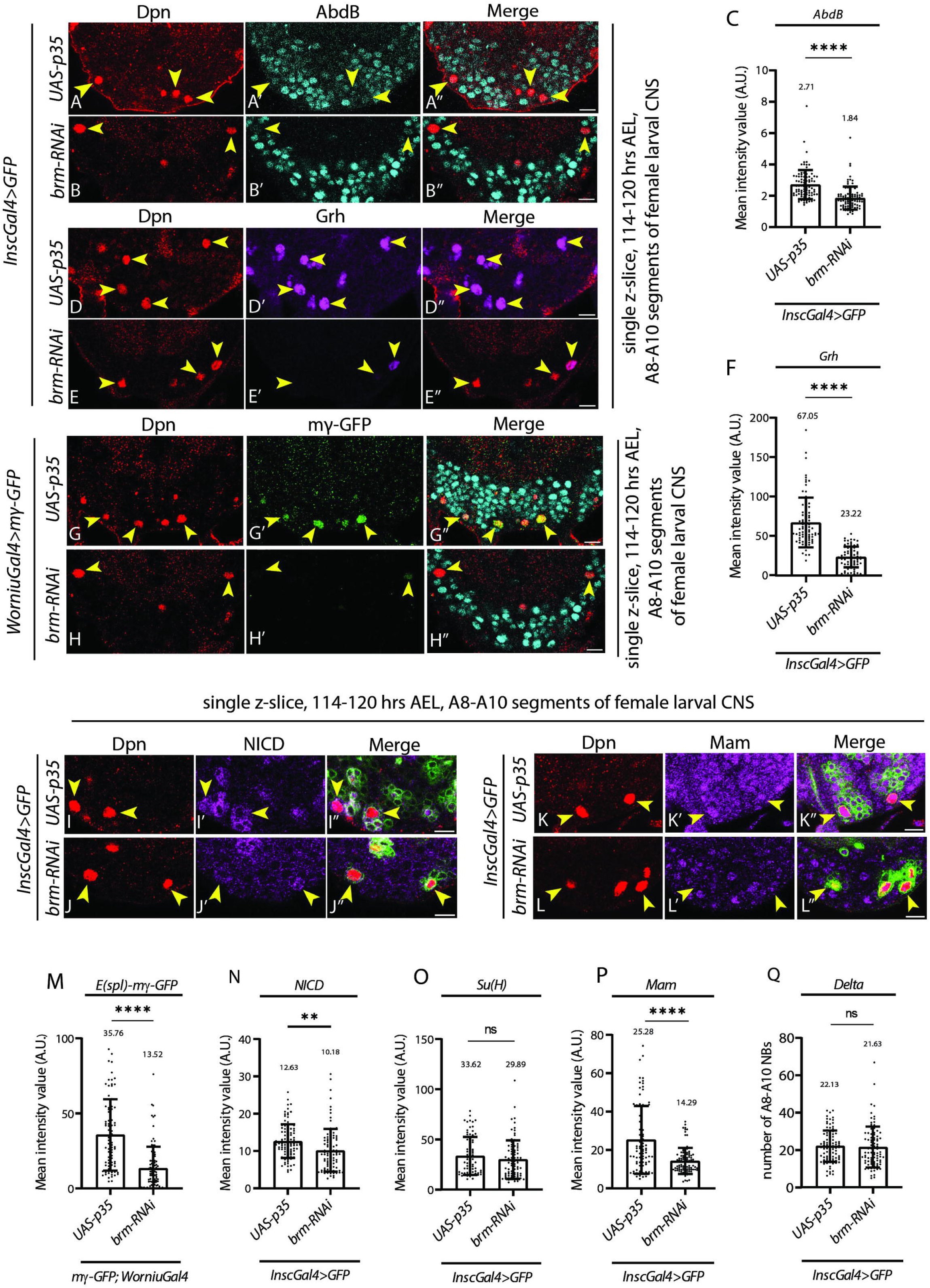
Brm controls A8-A10 NB apoptosis by regulating expression of Hox, Grh, NICD and Mam. (A-B, D-E, G-H, I-J, K-L) Show levels of Hox (Abd-B; 2.71±0.92; 93 NBs, *n*=9 VNCs *N*C:=C:3) (A), Grh (67.05±31.57; 79 NBs, *n*=7 VNCs *N*C:=C:3) (D), E(spl)-mγ-GFP (35.76±23.78; 78 NBs, *n*=10 VNCs *N*C:=C:5) (G), NICD (12.63±4.52; 99 NBs, *n*=8 VNCs N□=C:3) and Mam (25.28±17.69; 78 NBs, *n*=7 VNCs *N*C:=C:3) in control CNS, where NB apoptosis is blocked by expression of p35. Compared to these controls the knockdown of *brm* (*brm-RNAi*) leads to a significant reduction in the levels of Hox (1.84±0.73; 91 NBs, *n*C:=C:14 VNCs, *N*C:=C:3) (B), Grh (23.22±13.15; 73 NBs, *n*=10 VNCs, *N*C:=C:3) (E), and E(spl)-mγ-GFP (13.52±14.46; 97 NBs, *n*=10 VNCs *N*C:=C:5) (H), NICD (10.18±5.72; 95 NBs, n=14 VNCs, N=C:3) (J) and Mam (14.29±6.80; 94 NBs, *n*=13 VNCs *N*C:=C:3) (L). Graphs comparing the levels of Abd-B **(C)**, Grh **(F)**, E(spl)-mγ-GFP **(M)**, NICD **(N)**, Su(H) **(O)** and Mam **(P)**, Delta **(Q)** in A8-A10 NBs in p35 expressing control CNS and those with Brm knockdown (*brm-RNAi*). Yellow arrowheads indicate the ectopic NBs. Scale bars are 10 µm. Graph shows mean±SD. Significance is from a two-tailed Students’ unpaired *t-test*.

Thereafter, we tested if *brm* knockdown affected Notch signalling by regulating the expression levels of the members of the Notch signalling pathway. We checked the levels of the Notch receptor, N; the executive TF of Notch pathway, Su(H); ligand Delta, Dl; and co-activator, Mastermind (Mam). We observed that while levels of the Su(H) and Dl were unaffected (Fig. 2O and 2Q), the levels of NICD (Fig. 2J and 2N), and the co-activator Mam (Fig. 2L and 2P) were found to be significantly reduced in the NBs of A8-A10 segments.

These results collectively suggested that Brm knockdown blocks A8-A10 NB apoptosis by regulating Hox/Abd-B, Grh and Notch activity. The impact on the Notch activity seems to be through regulation of the levels of NICD, and Mam.

### Notch and Brm interact to regulate NB apoptosis

Brm is known to facilitate Su(H) binding on Notch target enhancers [36]. Considering this, we decided to check the interaction between Brm and Notch. We observed a weak but robust dominant interaction between the heterozygous null alleles of *brm* and *Notch* (*N^55e11/+^; brm^2/+^*), which showed a block of apoptosis of NBs in A8-A10 segments (Fig. 3D and 3E), this could be due to changes in the expression levels of Brm/Notch target genes or Brm’s interaction with the transcriptional complex. This was further substantiated by the double knockdown of Su(H) and Brm, wherein we observed a cumulative increase in the number of surviving NBs in A8-A10 segments (Fig. 3F), suggesting that the two are functioning together. The single knockdown of Su(H) also blocked NB apoptosis (Fig. 3F) and compromised Notch signalling (Fig. S2).

**Figure 3:**
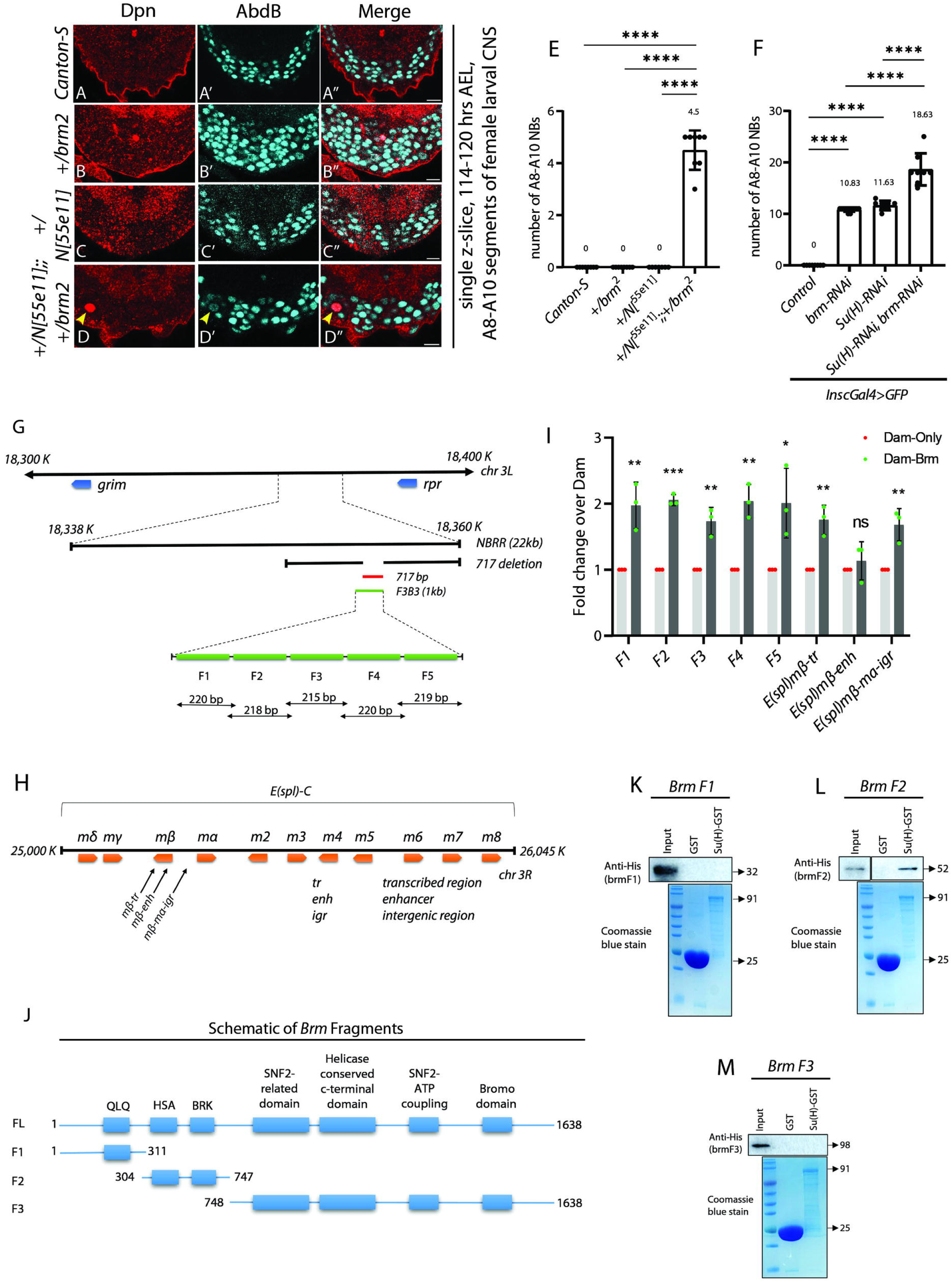
Brm binds to the *F3B3* apoptotic enhancer and dominantly interacts with the Notch signalling pathway. (A-D) Shows dominant interaction of *N^[55e11]^* and *brm^2^*. The terminal segments of the late L3 stage CNS show no surviving NBs in the case of control (*Canton-S;* 0±0 NBs, *n*C:=C:8 VNCs, *N*C:=C:2), *brm* heterozygotes (*brm^2^/+*; 0±0 NBs, *n*C:=C:8 VNCs, *N*C:=C:2), *Notch* heterozygotes (*N^[55e11]^/+*; 0±0 NBs, *n*C:=C:7 VNCs, *N*C:=C:3), but the double heterozygotes for *Notch* and *brm* (*N^[55e11]^/+*; *brm^2^/+*) shows ectopic NBs (4.50±0.75 NBs, *n*C:=C:8 VNCs, *N*C:=C:4). **(E)** Graph showing number of surviving NBs in A8-A10 segments in control (*CantonS*) and test (*brm^2^,* and *N^[55e11]^* heterozygotes and double heterozygotes *N^[55e11]^/+;brm^2^/+*) in late L3 stage CNS. **(F)** Graph showing number of surviving NBs in A8-A10 segments in controls (*inscGAL4>GFP*; 0±0 NBs, *n*C:=C:8 VNCs, *N*C:=C:3) and *brm-RNAi* (10.83±0.13 NBs, *n*C:=C:12 VNCs, *N*C:=C:3), *Su(H)-RNAi* (11.63±0.91 NBs, *n*C:=C:9 VNCs, *N*C:=C:3) and double knockdown (*brm-RNAi* and *Su(H)-RNAi;* 18.63±3.11 NBs, *n*C:=C:8 VNCs, *N*C:=C:4) in late L3 stage CNS. **(G)** Schematic showing the genomic location of *NBRR* region, *grim* and *reaper* genes on chromosome 3L. Green line shows the *F3B3* enhancer and red lines shows the extent of deletion. The 1 kb region is subdivided into 5 fragments of approximately ∼220 bp for *brm* binding study. **(H)** Schematic shows the representation of the genomic locus of *E(spl)-C* genes on Chr 3R. The regions indicated by arrow are the regions used as positive and negative controls for Brm binding by an earlier study (Pillidge and Bray, 2019). (**I**) Targeted DamID experiments show that Brm binds to the *F3B3* enhancer in the NBs. Fold enrichment graph of different fragments of *F3B3* for Dam-Brm compared to Dam-only (control) shows Brm binding onto the *F3B3* enhancer. *m*β*-tr* and *m*β*-m*α*-igr* are used as the positive controls for the binding and *m*β*-enh* as negative control. **(J)** Schematic showing domains of Brm protein (Brm-FL), and domains retained in deletion fragments of Brm, which are used for the in vitro pull-down. **(K-M)** Shows that Brm physically interacts with Su(H). GST tagged Su(H) could pull down BrmF2 **(L)**, but not BrmF1 **(K)** and BrmF3 **(M)**. Coomassie blue gel depicts the loading of GST-tagged Su(H). Yellow arrowheads indicate the ectopic NBs. Scale bars are 10 µm. Graph shows mean±SD. For zero-only control and non-zero test, significance is from Fisher’s test and for non-zero test and control, significance is from a Generalized Linear Model (GLM) with a binomial logistic regression performed using *‘R’.* For TaDa experiments with two independent groups, two-tailed unpaired student’s *t*-tests were used to test for significant differences.

A 1Kb *enhancer* (*F3B3*) (Fig. 3G), which starts expressing in the NBs in the mid-second instar stage, is suggested to be responsible for the activation of apoptotic genes *grim and reaper* and consequent death of A8-A10 NBs [22]. Since deletion of 717 bp of this *enhancer* blocks the NBs apoptosis (Δ*717/*Δ*717*) [22], we checked if Brm bound to this enhancer. For this, we used the Targeted DamID (TaDa) technique [43] to check for Brm binding onto the apoptotic enhancer in NBs of larval CNS. TaDa allows probing the tissue-specific in vivo binding status of chromatin/DNA binding proteins. To monitor the Brm binding onto the 1 Kb apoptotic enhancer, we divided it into five fragments of approximately 200 bps and designed primers for these amplicons to assay the enrichment of Brm-bound fragments using RealTime (RT) PCR (Fig. 3G). Using the TaDa-RT PCR, we found that Dam-Brm fusion showed approximately 2-fold enrichment compared to the Dam-only control for all five fragments (Fig. 3I), suggesting that Brm bound to the 1 Kb apoptotic enhancer in the NBs. We could also observe the Brm binding onto the intergenic region between mβ and mα genes and mβ-tr (mβ-transcribed region) of the Enhancer of Split [E(spl)] complex (Fig 3H and 3I). E(spl) complex is a known target for Notch signalling, and the intergenic region between E(spl)-mβ and mα is known to be bound by Brm [36], which therefore acted as positive controls and E(spl)mβ-enh acted as a negative control. Subsequently, we reanalysed an previously published Targeted DamID-seq data [44] for NBs in the early L3 stage of development and found that Brm bound to the 1kb enhancer (*F3B3*) in NBs (Fig. S3), reaffirming our results. In TaDa-Seq data we also observed a significant Brm binding on apoptotic genes, *grim* and *reaper*, in addition to *N* and *Mam* (Fig. S3).

Considering our genetic data (dominant genetic and RNAi interactions) as well as previously published results, where Brm binding is required for Su(H) to bind to DNA [36], we expected that Brm and Su(H) may physically interact with each other. Therefore, we checked this using an in vitro pulldown experiment wherein different regions of Brm (Fig. 3J) were tested for their interaction with Su(H). Here we found that Brm and Su(H) interact with each other through HSA and BRK domains of Brm (Fig. 3J-3M), which are important for Brm’s interaction with chromatin and act as a primary binding platform for actin and ARP (actin related proteins) to regulate chromatin remodelling function [45–47].

Collectively, these results show that Brm and Notch exhibit a dominant genetic interaction, with Brm binding to the enhancer which may be required for NB apoptosis. We also find that Brm physically interacts with Su(H) through its HSA and BRK domains.

### Brm regulates *grim* and *reaper* genes through the *F3B3* enhancer

Next, we functionally tested whether the *brm* knockdown-mediated block of NB apoptosis happened through its impact on the *F3B3* enhancer. For this, we checked the expression of *F3B3-lacZ* in *brm* knockdown as well as in the case of overexpression of the catalytically dead version of Brm (Brm-K804R). The controls in this case were the larvae, where NB apoptosis in the CNS was blocked by expression of the cell death blocker p35. Here, we observed a significant decrease in the percentage of cells expressing *F3B3-lacZ* in *brm* knockdown and overexpression of Brm-K804R compared to NBs in p35 expressing control CNS (Fig. 4A). We also tested knockdown for Notch and Mam for their impact on *F3B3-lacZ* line expression in NBs. The control VNCs typically have 70% of the A8-A10 NBs expressing *F3B3-lacZ* in the early L3 stage; this number was reduced to 26% in the case of *brm*-knockdown, 32% in case of overexpression of BrmK804R and 17% and 8% in the case of *Notch* and *mam* knockdown (Fig. 4A). We also observed that knockdown for *mam* resulted in block of NB apoptosis (Fig. S2E) and reduction in Notch activity (Fig. S2I). These results suggest that the block of apoptosis that is seen in the case of *brm, Notch* and *Mam* knockdowns, and in the case of the overexpression of catalytically dead Brm, is mediated through their impact on the *F3B3* enhancer. Next, we checked if the homozygous deletion of the apoptotic enhancer (Δ*717/*Δ*717*) resulted in downregulation of the *grim* and *reaper* transcripts in the CNS and observed an approximately a 50% decrease in the transcript levels (Fig. 4B).

**Figure 4:**
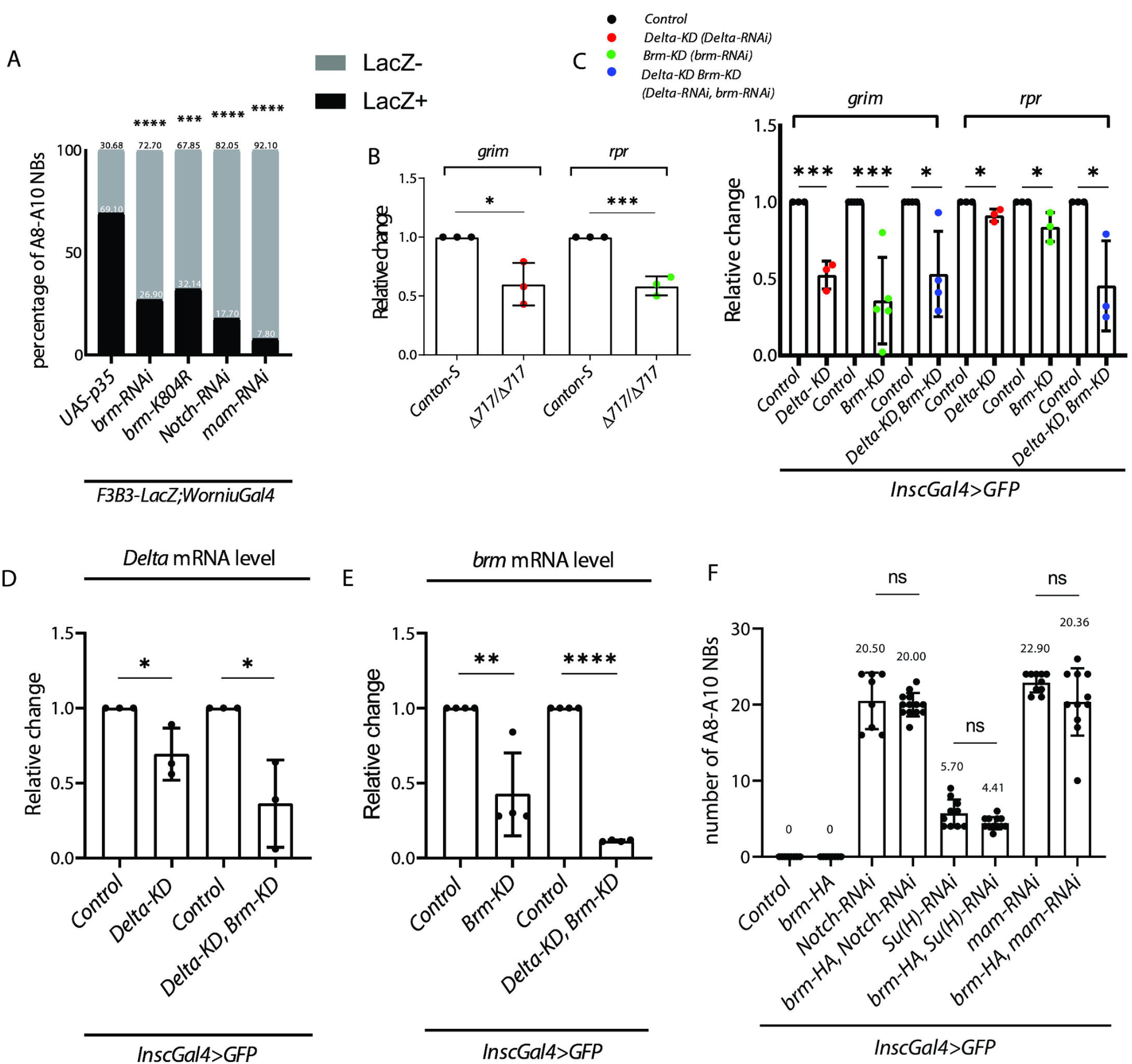
Brm regulates transcription of *grim* and *reaper* through the *F3B3* enhancer. **(A)** Graph comparing the percentage of LacZ*-*positive and LacZ-negative NBs in A8-A10 segments of in late L3 stage CNS; p35 expressing control larvae are compared with *brm-RNAi, brm-K804R, Notch-RNAi* and *Mam-RNAi.* **(B)** Graph shows that compared to controls (*Canton S*) the transcript levels of *grim* and *rpr* are reduced in homozygous deletion for apoptotic enhancer *F3B3* (Δ*717/*Δ*717)*. The result suggests that *brm* regulate transcripts of *grim* and *reaper* through *F3B3* apoptotic enhancer. **(C)** Graph compares the transcript levels of *grim* and *reaper* in *Dl-RNAi* (*Delta*-KD), *brm-RNAi* (*brm-KD*) and collectively in double knockdown for the two (*Delta-KD+brm-KD*). The transcript levels for *grim* are reduced in single knockdown; the extent of reduction in double knockdown is similar to single *Delta-KD*, suggesting that Notch is downstream of Brm. **(D-E)** Shows that the transcript levels of *Dl* and *brm* go down in the individual and double knockdowns. **(F)** Graphs show the number of surviving NBs in A8-A10 segments for the controls (0±0 NBs, *n*C:=C:8 VNCs, *N*C:=C:3), *brm* overexpression (0±0 NBs, *n*C:=C:10 VNCs, *N*C:=C:3), *Notch-RNAi* (20.50±3.70 NBs, *n*C:=C:8 VNCs, *N*C:=C:3), *Su(H)-RNAi* (5.70±1.82 NBs, *n*C:=C:10 VNCs, *N*C:=C:3), *mam-RNAi* (22.90±1.28 NBs, *n*C:=C:10 VNCs, *N*C:=C:4) and the rescue experiments where *brm* is over-expressed in *Notch-RNAi* (20.00±1.52 NBs, *n*C:=C:13 VNCs, *N*C:=C:3), *Su(H)-RNAi* (4.41±0.79 NBs, *n*C:=C:12 VNCs, *N*C:=C:3) and *mam-RNAi* (20.36±4.43 NBs, *n*C:=C:11 VNCs, *N*C:=C:4) backgrounds. These experiments establish that *Notch* is downstream of *brm,* since *brm* overexpression fails to rescue the knockdowns of *Notch, Su(H) and Mam.* Graph indicating mean±SD. The transcript levels in panels **B**, **C**, **D** and **E** are normalised to GAPDH expression levels in control and tests. For zero-only control and non-zero test, significance is from Fisher’s test and for non-zero test and control, significance is from a Generalized Linear Model (GLM) with a binomial logistic regression performed using *‘R’.* For experiments with two independent groups, two-tailed unpaired student’s *t*-tests were used to test for significant differences.

Considering the reduction in the number of *F3B3-positive* NBs in *brm* knockdown, we decided to check the impact of the blocking Notch signaling and *brm*-Knockdown conditions individually and collectively on the *grim and reaper* transcript levels. For this, we did individual knockdown of Notch ligand *Delta* (*Delta*-KD) and *brm* gene (*brm*-KD), and a combined knockdown of the two. In the case of *reaper* gene, we observed only a modest decrease in transcript levels in case of *Delta*-KD and *brm*-KD. However, in the case of *grim* gene, we observed a 50% and 60% decrease in the transcript levels in the case of *Delta*-KD and *brm*-KD conditions, respectively (Fig. 4C). The reduction in *Delta* and *brm* transcripts in response to RNAi mediated knockdown was confirmed in the case of single (*Delta*-KD, *brm*-KD) and collective knockdowns (Fig. 4D and 4E).

In the case of double knockdown, we observed only a 50% decrease in *grim*, similar to what is seen in the case of *Delta*-KD condition, which suggested that Notch signalling functions downstream of the Brm to regulate apoptotic genes. To probe the hierarchical relation between Notch and Brm, we also assayed the number of surviving A8-A10 NBs (Fig. 4F). To this end, we overexpressed Brm in the case of Notch, Su(H) and Mam knockdowns (which blocked NB apoptosis to varying extents). In all these cases, we observed that overexpression of Brm did not result in any change in the block of NB apoptosis seen in A8-A10 segments, thereby further establishing that Brm is genetically upstream to Notch signalling pathway (Fig. 4F).

Considering these results and Targeted DamID data, we favour the idea that Brm regulates apoptosis by modulating the expression of *grim* and *reaper* genes, through its direct binding to the *F3B3* enhancer. Collectively, our results suggest that Brm functions upstream of the Notch signalling pathway, but also collaborates with it at the level of the *F3B3* enhancer.

### Drosophila CBP/p300 recruitment onto the apoptotic enhancer is important for Notch activity and apoptosis

A rapid and large-scale change in the H3K56ac is reported in case of Su(H) bound Notch target enhancers upon Notch signalling activation. This is dependent on CBP/p300 (encoded by the *nejire* (*nej*) gene in *Drosophila*) and leads to increased transcription following Notch signalling activation [35]. Therefore, we wanted to test the role of Nej in A8-A10 NB apoptosis. Since, the H3K56ac antibody did not show good staining in NBs, we monitored H3K27ac of the nucleosomes (also mediated by Nej) and to assess its role in NB apoptosis (Tie et al, 2012).

We observed that *nej* knockdown resulted in the block of NB apoptosis in A8-A10 segments (Fig. 5B and 5L). Expectedly, we also observed that levels of H3K27ac in the surviving A8-A10 NBs were reduced in the case of the *nej* knockdown (Fig. 5D’ and 5M). Considering that Nej mediated histone acetylation is important for increased transcription of the Notch targets, we checked the E(spl)mγ-GFP expression in these NBs and observed a significant reduction in its levels in the case of *nej* knockdown (Fig. 5F’ and 5N). Next, we checked the impact of *nej* knockdown on the apoptotic enhancer. We observed a significant decrease in the percentage of cells expressing *F3B3-lacZ* compared to controls (Fig. 5G), suggesting that the block of apoptosis in *nej* knockdown was mediated through its impact on the enhancer. Thereafter, we checked if Brm knockdown affected Nej levels in the NBs. Here, we observed a 40% reduction in the levels of Nej (Fig. 5I and 5O) and a decrease in H3K27ac marks on the nucleosomes in the surviving A8-A10 NBs (Fig. 5J’ and 5M). These results suggested that *brm* is upstream of *nej*, which was further supported by the fact that its overexpression did not rescue the block of NB apoptosis and the reduction in the levels of H3K27ac seen in the case of *nej* knockdown (Fig. 5K’, 5L and 5M). Congruent to this, we could observe significant Brm binding on the *nej* gene (Fig S3) in published TaDa-Seq data for Brm [44].

**Figure 5:**
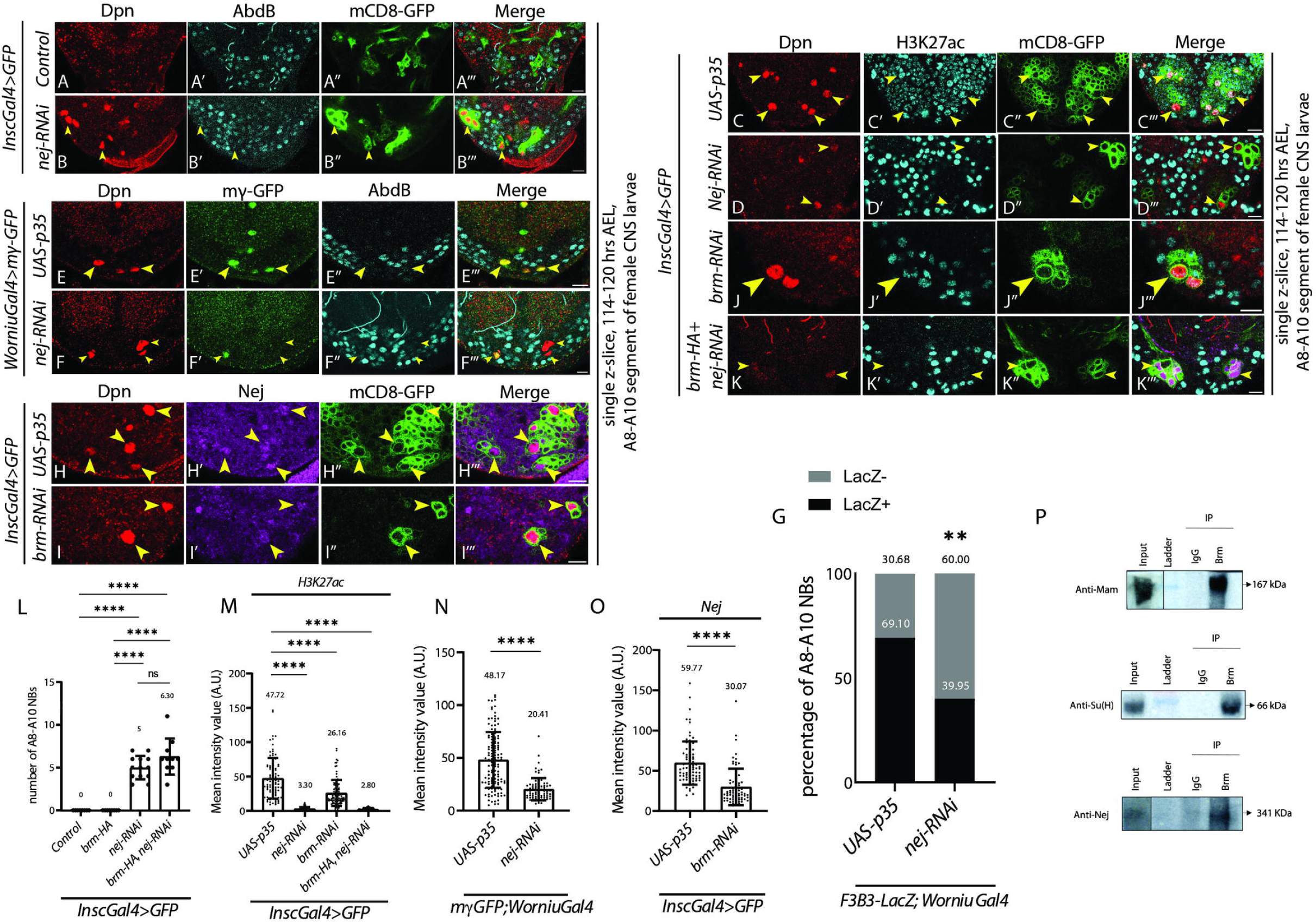
***Drosophila* ortholog of histone acetyl transferase CBP/p300 (Nej) is required for NB apoptosis and is regulated by *brm*. (A-B)** Compared to controls (*inscGAL4>GFP*; 0±0 NBs, *n*C:=C:9 VNCs, *N*C:=C:3) knockdown of *Drosophila* CBP/p300 (*nej-RNAi*; 5.00±1.35 NBs, *n*C:=C:13 VNCs, *N*C:=C:3) blocks A8-A10 NB apoptosis. **(E-F)** The knockdown of Nej (*nej-RNAi*) leads to a significant reduction in the levels of E(spl)-mγ-GFP (20.41±10.58, 81 NBs, *n*=9 VNCs, *N*C:=C:3) (F) in A8-A10 NBs compared p35 expressing control NBs (48.17±26.34, 131 NBs, *n*=7 VNCs, *N*C:=C:2) (E) (blocked for apoptosis); indicating that Notch activity is abrogated in Nej knockdown. **(H-I)** The knockdown of *brm* (*brm-RNAi*) leads to a significant reduction in the levels of Nej (30.07±22.68, 76 NBs, *n*=10 VNCs *N*C:=C:2) (I) in A8-A10 NBs compared to p35 expressing control NBs (59.77±26.90, 78 NBs, *n*=7 VNCs *N*C:=C:2) (H), indicating that Nej levels are affected in the case of *brm* knockdown. **(C-D, J-K)** Shows a significant reduction in the levels of H3K27ac in the case of the knockdown of Nej (*nej-RNAi;* 3.3±2.37, 112 NBs, *n*=11 VNCs, *N*C:=C:3) (D) and *brm* (*brm-RNAi;* 26.16±18.78, 97 NBs, *n*=17 VNCs, *N*C:=C:4) (**J**) in A8-A10 NBs compared to p35 expressing control (47.72±29.46, 93 NBs, *n*=6 VNCs, *N*C:=C:2) (C). The reduction in H3K27ac levels seen in the case of Nej knockdown is not rescued by overexpression of *brm* (2.80±1.36, 91 NBs, *n*=9 VNCs *N*C:=C:2) in these cells, indicating that Nej is downstream of Brm. **(G)** Graph comparing the percentage of LacZ*-*positive and LacZ-negative NBs in A8-A10 segments of in late L3 stage CNS; p35 expressing control larvae are compared with *nej-RNAi.* The result suggests that *nej* impacts NB apoptosis through *F3B3* enhancer. **(L)** Graph comparing the number of surviving A8-A10 NBs in controls (*inscGAL4>GFP*; 0±0 NBs, *n*C:=C:8 VNCs, *N*C:=C:2), *brm* overexpression (0±0 NBs, *n*C:=10 VNCs, *N*C:=C:3), *nej* knockdown (*nej-RNAi*; 5.00±1.35 NBs, *n*C:=C:13 VNCs, *N*C:=C:3) and *brm* overexpression in the background of *nej* knockdown (6.30±2.11 NBs, *n*C:=C:10 VNCs, *N*C:=C:5). **(M)** Shows the graph comparing the levels of *H3K27ac* level in A8-A10 NBs expressing *p35, nej-RNAi, brm-RNAi and brm* overexpression in *nej-RNAi*. The data in (L) and (M) show that Brm overexpression could not rescue Nej knockdown phenotype, establishing that Nej functions downstream of Brm. **(N)** Shows that Nej knockdown reduces E(spl)-mγ-GFP expression levels in A8-A10 NBs compared to p35 expressing control NBs (blocked for apoptosis). **(O)** Shows that Brm knockdown results in downregulation of Nej levels in A8-A10 NBs compared to p35 expressing control NBs (blocked for apoptosis). **(P)** Western blot showing that Brm could pull down Su(H), Mam and Nej from larval lysates. IgG in the first lane is used as a negative control, which does not interact with components pulled down in the experiment. Yellow arrowheads indicate the ectopic NBs. Scale bars are 10 µm. Graph shows mean±SD. For zero only control and non-zero test, significance is from Fisher’s test and for non-zero test and control, significance is from a Generalized Linear Model (GLM) with a binomial logistic regression performed using *‘R’.* For experiments with two independent groups, two-tailed unpaired student’s *t*-tests were used to test for significant differences. A one-way ANOVA and a subsequent Dunnett’s post doc test was used to compare multiple hypothesis.

Since our previous results supported the idea that one of the ways Brm mediates the block of NB apoptosis is by binding to the *F3B3* enhancer, where it may interact with Su(H). We speculated that upon signalling activation in presence of Grh and Abd-B, NICD binds to the signalling complex (Su(H) and Mam) and recruits Nej to increase acetylation at the *F3B3* enhancer. If this were to be the case, we expected that all these factors might function as a complex in vivo. Congruent with this, Brm could pull down Su(H), Mam and Nej from larval lysates (Fig. 5P).

Our results suggest that Nej-mediated H3K27 acetylation at the Notch-sensitive *F3B3* enhancer contributes to NB apoptosis. The pulldown from whole larval lysates confirms that components of the Notch pathway, Brm, and Nej form a complex, supporting the idea that Brm interacts with the Notch-Nej complex to facilitate NB apoptosis.

## Discussion

SWI-SNF ATPase Brm (and its orthologs) and Notch signalling have been known to interact in different cellular contexts. During heart development in Zebrafish, Brg1 is known to interact with lysine demethylase to regulate Notch receptor expression [48]. Similarly, in T cell acute lymphoblastic leukemia cells, it has been shown that NICD-Su(H)-Mam orthologs along with PBAF complex and histone demethylases, promote epigenetic modification at Notch target genes [49]. In the context of Left-Right asymmetry, it has been shown that Baf60 knockdown fails to activate Notch target gene Nodal in vivo; thereafter, using cell culture, it was shown that Baf60 is required for Notch dependent transcriptional activation by stabilizing the interaction between activated Notch and RBPJ-1 (Su(H) ortholog) [50]. Similarly, in Drosophila also, Brm and Notch signalling have been shown to interact during development [36–38]. A dominant interaction between Notch ligand Dl and the ATPase dead version of Brm was reported previously [38]. More recent work in Drosophila cell culture showed that Brm facilitate Su(H) binding to Notch target loci [36–38]. Subsequent study in wing disc showed that the Brm-associated BAP complex (conventionally known for transcriptional activation) could execute gene repression as well, wherein Osa (a component of the BAP complex) bound to components and targets of the Notch signalling pathway, thereby constraining their expression [37]. Expectedly, the loss of Osa resulted in activation of the Notch ligand Delta in the wing disc. Some of these examples (mainly from vertebrate cell culture studies), showed a direct interaction between Brm/BAP and Notch/Su(H) complexes to regulate the Notch target loci similar to the role of the Brm in mediating the binding of Su(H) onto the DNA to facilitate Notch-dependent transcription in Drosophila [36]. However, the in vivo validation of this interaction in a physiological context has been missing. We have investigated the molecular mechanism of this interaction in the context of Hox-dependent NB apoptosis in larval CNS. Our results suggest a multi-tier regulation of NB apoptosis by SWI-SNF ATPase Brahma in the larval CNS. The first tier of regulation is in controlling the expression of the *Drosophila* homolog of CBP/p300 (Nej-a HAT) and the molecules directly involved in the cell death (Hox/Abd-B, Grh, and members of the Notch signalling pathway-N and Mam) (Fig 6A). The second tier of regulation is by direct binding by Brm and modulation of the chromatin accessibility at the apoptotic enhancer. This facilitates the binding of Su(H) to the enhancer thereby keeping it ready to respond to the apoptotic trigger (an increase in the Notch activity and the levels of Grh in presence of Abd-B) (Fig 6B). Our data support a model in which, in response to the apoptotic trigger, the Brm and Su(H)/Mam complex recruit CBP/p300 (Nej) onto the apoptotic enhancer, thereby increase the H3K27ac marks on the nucleosomes to further open up the chromatin, which in presence of Grh and Abd-B facilitates *grim* and *reaper* transcription in NBs of A8-A10 segments (Fig 6C). The role of Brm in regulating H3K27ac through its direct interaction with CBP has been reported earlier [51], though the Brm-CBP-mediated modulation of Notch signalling in the context of NB apoptosis is novel.

**Figure 6:**
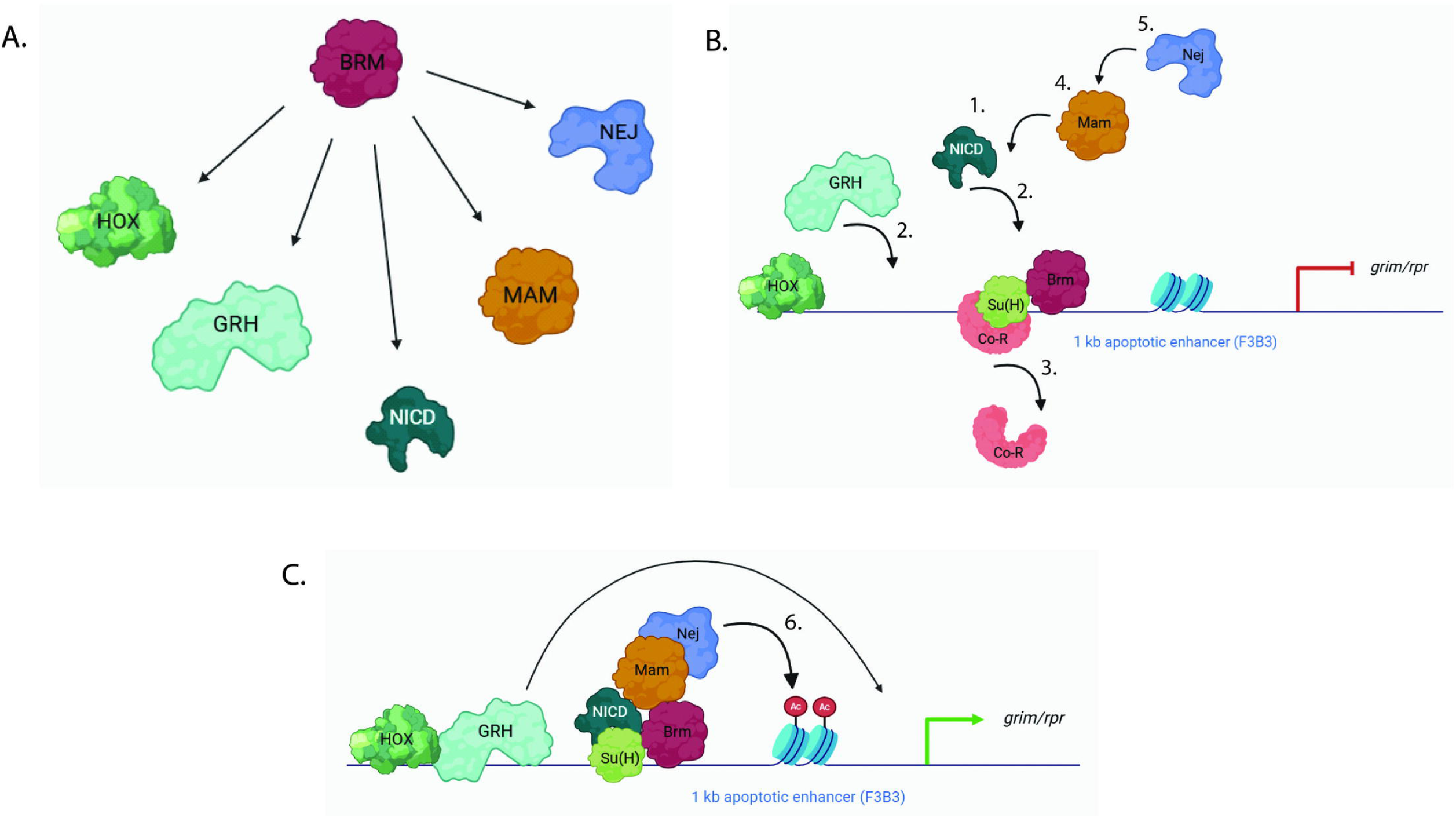
(**A**) Schematic suggesting that Brm regulates the levels of Hox (Abd-B), Grh, and components of Notch signalling pathway (NICD, Mam) and Histone Acetyl Transferase CBP/p300/Nej. **(B)** Prior to activation of Notch signalling, Brm opens the chromatin and facilitates the binding of Su(H) onto DNA. **(C)** Upon activation of Notch signalling and increase in Grh levels (in presence of Abd-B), NICD binds to Su(H) and the corepressor is removed and coactivator Mastermind is recruited, which then recruits histone acetyl transferase CBP/p300/Nej to the Notch signalling complex, increasing the H3K27ac, resulting in further opening of the chromatin and transcriptional activation of *grim* and *reaper*.

Considering the role of the Brm as a chromatin remodeller, we expected it to regulate NB cell death by positively regulating the expression of the *RHG* family of apoptotic genes. Congruent with this, the knockdown of Brm, Moira and Snr1, which are core components of the Brm complex and Osa, a BAP complex-specific subunit, blocked NB apoptosis (Fig. S1). The knockdown of BAP111 and BAP55, which are also part of the BAP complex, did not show an apoptosis block, we believe this may be because the RNAi knockdown was not potent enough to show a phenotype.

Furthermore, since the knockdown of any of the PBAP complex members like SAYP, BAP170, and Polybromo did not block of NB apoptosis (Fig. S1), while knockdown of Osa did, suggests that PBAP complex does not play a role in NB apoptosis thereby further reiterating the role of BAP complex in apoptotic gene activation.

The first tier of regulation mentioned above is one way of interaction between Brm and the Notch signalling pathway. Other supporting evidence is the dominant genetic interaction between mutants of *Notch* and *brm* heterozygotes, and the double knockdown of Brm and Su(H), both of which resulted in the block of NB apoptosis. This is expected considering a previous report where the Notch ligand Dl showed a dominant genetic interaction with the ATPase dead Brm [38]. However, our data also supports the idea that Notch-Brm interaction also happens at the level of the apoptotic enhancer. This is supported by the fact that both Su(H) [22] and Brm bind to the apoptotic enhancer (TaDa data), and their respective knockdowns (this study and [22]) reduced the expression of *F3B3-lacZ* in the NBs of the CNS. On these lines, we observed that individual knockdown of Brm did not affect Cut (a target of Notch signalling) expression in the NBs, but the double knockdown of Brm and Su(H) reduced the expression of Cut to an extent greater than what is seen for Su(H) knockdown alone (Fig. S5), further suggesting for a collective role of the two “on the DNA” rather than “off the DNA”. Other supporting evidence is the in vivo pulldown from the larval lysates, which show that Brm, Su(H), Mam, and CBP/Nej may exist as a complex in a cell, which is likely to be on the DNA, considering that all of them are DNA/chromatin binding proteins. The in vitro pulldown showing that Brm and Su(H) physically interact with each other through Brm’s highly conserved HSA and BRK domains, lead us to further favour the idea that this interaction may be happening on the apoptotic enhancer, since these domains act as a binding platform for actin-related proteins (ARPs) and help Brm to perform its remodelling function [45–47]. Lastly, whether disrupting this interaction can actually impact the in vivo functionality of the complex and block Hox-dependent NB apoptosis needs to be tested.

## Supplementary Figures

**Fig. S1: BAP Complex is responsible for NB apoptosis** (A) Schematic showing the components of BAP and PBAP complexes. The core subunits are shown in 3 colors: pink (Moira, Snr1), green (Bap111, Bap60, Bap55 and Actin) and red (Brm). The complex-specific subunits are blue (Osa for BAP; Polybromo, BAP170, Sayp for PBAP). (B-I) Knockdown of Snr1 and Moira results in a block of A8-A10 NB apoptosis. The CNS from controls (*inscGAL4>GFP*; 0±0 NBs, *n*C:=C:8 VNCs, *N*C:=C:2) (B) show no surviving NBs while knockdown for Moira (*Moira-RNAi-1 and 2;* 5.91±1.67 NBs, *n*C:=C:18 VNCs, *N*C:=C:6; 7.57±1.83 NBs, *n*C:=C:9 VNCs, *N*C:=C:3) (C, D) and Snr 1 (*Snr1-RNAi-1 and 2;* 8.42±2.63 NBs, *n*C:=C:15 VNCs, *N*C:=C:4; 5.50±3.39 NBs, *n*C:=C:7 VNCs, *N*C:=C:4) (E, F) blocks NB apoptosis in late L3 CNS. Knockdown for Bap111 (G-H) and Bap55 (I) did not show any surviving NBs. **(J-K)** Compared to controls (*inscGAL4>GFP*; 0±0 NBs, *n*C:=C:8 VNCs, *N*C:=C:2) (B), knockdown of BAP complex specific subunit Osa (*Osa-RNAi-1 and 2;* 6.53±1.71 NBs, *n*C:=C:15 VNCs, *N*C:=C:3, 5.66±2.29 NBs, *n*C:=C:10 VNCs, *N*C:=C:4) (J-K) results in a block of A8-A10 NB apoptosis. In contrast, the knockdown of PBAP-specific subunits Sayp, Polybromo and Bap170 (L-Q) does not block NB apoptosis. (R-T) Graph showing the surviving A8-A10 NBs in case of the knockdown for core (R) and BAP (S) and PBAP (T) specific subunits of the complex. (U-U’’’ and V-V”’) Shows *brm-RNAi* rescue with the human ortholog of Brm (hSMARCA2). (U-U’’’) Over-expression of HA-hSMARCA2 in the A8-A10 region shows no block of apoptosis (0.00±0.49 NBs, *n* = 10 VNCs, *N* = 3). (V-V’’’) Over-expression of HA-hSMARCA2 could rescue (3.07±0.49 NBs, *n* = 14 VNCs, *N* = 4) *brm-RNAi phenotype* (10.83±0.13 NBs, *n* = 12 VNCs, N = 3). Yellow arrowheads indicate the ectopic NBs. Scale bars are 10 µm. Graph shows mean±SD. For zero-only control and non-zero test, significance is from Fisher’s test and for non-zero test and control, significance is from a Generalized Linear Model (GLM) with a binomial logistic regression performed using *‘R’*.

**Fig. S2: Knockdown of *Notch, Su(H) and Mam*** (A-D) Knockdown of Notch, Su(H) and Mam results in a block of A8-A10 NB apoptosis. The CNS from controls (*inscGAL4>GFP*; 0±0 NBs, *n*C:=C:8 VNCs, *N*C:=C:2) (A) show no surviving NBs while knockdown for Notch (*Notch-RNAi-1* 21.50±5.01 NBs, *n*C:=C:8 VNCs, *N*C:=C:3) (B), Su(H) (*Su(H)-RNAi;* 11.63±0.30 NBs, *n*C:=C:8 VNCs, *N*C:=C:3) (C), and Mam (*mam-RNAi*; 24.10±0.57 NBs, *n*C:=C:10 VNCs, *N*C:=C:3) (D) blocks NB apoptosis in late L3 CNS. (F-H) Show levels of E(spl)-mγ-GFP in control NBs blocked for apoptosis by expression of p35 (48.17±26.34, 131 NBs, *n*=7 VNCs *N*C:=C:3) (F). The E(spl)-mγ-GFP expression is significantly reduced in A8-A10 NBs upon the knockdown of Su(H) (*Su(H)-RNAi*) (G) and Mam (*mam-RNAi*) (H) in the late L3 stage. (E) Graph showing the number of surviving A8-A10 NBs in late L3 stage CNS in *Control (inscGAL4>GFP), Notch-RNAi, Su(H)-RNAi* and *Mam-RNAi*. (I) Graph comparing the levels of E(spl)-mγ-GFP in A8-A10 NBs of controls, Su(H) and Mam knockdown. Yellow arrowheads indicate the ectopic NBs. Scale bars are 10 µm. Graph shows mean±SD. For zero-only control and non-zero test, significance is from Fisher’s test and for non-zero test and control. For experiments with two independent groups, two-tailed unpaired student’s *t*-tests were used to test for significant differences using GraphPad prism 8.0. A one-way ANOVA and a subsequent Dunnett’s post doc test was used to compare multiple hypothesis.

**Fig. S3: Targeted Dam ID shows that Brm binds to *grim, reaper, F3B3, Notch, mam, nej* and *Su(H)*** Shows binding profile of Brm on *grim (A), reaper (A)*, *F3B3* apoptotic enhancer (A), *Notch* (B), *mam* (C), and *nej* (D) in NBs at the mid L3 stage in larval CNS as identified by Targeted Dam ID. The binding profiles suggest that Brm binds and regulates these genes. Y-axis limits are set from −10 to 10 for occupancy comparison across the tracks. The coordinates indicated at the top are in kilobases.

**Fig. S4: Temperature shift protocols** (A-C) Show the temperature shift (TS) protocols used in different experiments. Blue lines indicate when the flies were grown at 18°C. Red lines indicate the time for which the flies were grown at 29°C. Black line indicate the developmental timeline for at 25°C. The black downward facing arrow indicates the time of dissection of the larvae. The L3 stage has been divided into three 16 hr intervals to define the early, mid, and late L3 stage.

**Fig. S5: Double knockdown of *Su(H)* and *brm* reduces Cut levels in A8-A10 NBs** (A-D) Show levels of Cut (10.16±3.99, 106 NBs, *n*=7 VNCs *N*C:=C:3) (A), in control A8-A10 NBs blocked for apoptosis by expression of p35. The Cut expression is significantly reduced in A8-A10 NBs upon the knockdown of Su(H) (*Su(H)-RNAi;* 5.17±2.39, 91 NBs, *n*=8 VNCs *N*C:=C:3) (B) and double knockdown for Su(H) and Brm (*Su(H)-RNAi* and *brm-RNAi*; 3.39±1.08, 107 NBs, *n*C:=C:8 VNCs, *N*C:=C:3) (D), but not in case of Brm knockdown alone (*brm-RNAi*; 9.74±3.51, 101 NBs, *n*C:=C:15 VNCs, *N*C:=C:3) (C), in the late L3 stage. (E) Graph comparing the levels of Cut in control, single knockdown for *Brm* and Su(H); and double knockdown for the two together. Yellow arrowheads indicate the ectopic NBs. Scale bars are 10 µm. Graph shows mean±SD. A one-way ANOVA and a subsequent Bonferroni’s post doc test was used to compare multiple hypothesis.

**Fig. S6: Expression of *F3B3-lacZ* in terminal NBs upon over-expressing Brm-K804R, and knockdown of *brm*, *nej* and components of Notch pathway.** (A’ B’ C’ D’ E’ and F’) Shows *F3B3-lacZ*+ vs *F3B3-lacZ-* NBs from the representative images of terminal (A8-A10) segments of late L3 stage female brain of control over-expressing p35 (A-A”’), *brm-RNAi* (B-B’’’), over-expression of Brm-K804R (C-C’’’), *Notch-RNAi* (D-D’’’), *mam-RNAi* (E-E’’’) and *nej-RNAi* (F-F’’’). Yellow arrowheads indicate the ectopic NBs. Scale bars are 10 μm.

**Fig. S7: Brm’s interaction with Nej and components of Notch pathway Su(H), Mam.** Uncropped pictures of the western blot showing that Brm could pull down (A) Su(H), (B-C) Mam and (D-E) Nej from larval lysates.

## Materials and Methods

### Statistical analysis

For the count-based experiments with a control group with zero values and a test with non-zero values, we conducted Fisher’s exact test using GraphPad Prism 8.0. And for the count-based data with non-zero control and test values, we performed a generalised linear regression (GLM) with a Binomial Logistic regression to determine the differences using “R”. For experiments with two independent groups, two-tailed unpaired Student’s *t*-tests were used to test for significant differences using GraphPad prism 8.0. For experiments with more than two genotypes with a single control, significant differences amongst specific genotypes were tested using a one-way ANOVA and a subsequent Dunnett’s post hoc test. A one-way ANOVA and a subsequent Bonferroni’s post doc test was used to compare multiple hypothesis with control and within groups as well. For all graphs, error bars represent SD. *P* and Adjusted-*P* values are reported as follows: *P*C:>C:0.05, ns (not significant); \**P*C:<C:0.05; \*\**P*C:<C:0.01; \*\*\**P*C:<C:0.001; \*\*\*\**P*C:<C:0.0001.

### Fly stocks

Following lines were used: *Canton-S* (BDSC 64349), *brm-RNAi* (BDSC 31712, VDRC 37720, VDRC 37721), *Delta*-RNAi (BDSC 36784), *Notch*-RNAi (BDSC 28981), *Su(H)*-RNAi (BDSC 28900, VDRC 103597), *nej-RNAi* (BDSC 27724, BDSC 37489), *UAS-HA-hSMARCA2* (BDSC 84943), *N^[55e11]^* (BDSC 28813), *brm^d00415^* (BDSC 19144), *brm^[I21]^* (BDSC 4108), *brm^2^* (BDSC 3619), *brm^K804R^* (Well Genetics for this study). *UAS-HA-brm* (this study), *UAS-HA-brm*-K804R (this study), *UAS-p35* (DGRC 108019), *UAS-dcr2; inscGAL4 UASmCD8-GFP and UAS-dcr2; inscGAL4 UASmCD8-GFP; tub-GAL80^ts^ (*J. A. Knoblich), *worniu-GAL4* (C. Doe), *UAS-LT3-Dam, UAS-LT3-Dam-Brm* (A. Brand [44]), *E(spl)m*γ*-GFP* (S. Bray [42]), *F3B3-LacZ* [20], Δ*717* [22].

CRISPR-mediated mutagenesis for *brm^K804R^* allele was performed by WellGenetics Inc. using modified methods of Kondo and Ueda [52]. In brief, gRNA sequence AAGGTAGGTTACCAGCGAAA[TGG] were cloned into U6 promoter plasmid(s). Cassette PBacDsRed, which contains 3xP3-DsRed flanked by PiggyBac terminal repeats, and two homology arms with point mutation were cloned into pUC57-Kan as donor template for repair. *brm*-targeting gRNAs and hs-Cas9 were supplied in DNA plasmids, together with donor plasmid for microinjection into embryos of control strain *w^[1118]^*. F1 flies carrying selection marker of 3xP3-DsRed were further validated by genomic PCR and sequencing. CRISPR generates a break in *brm* and is replaced by cassette PBacDsRed with the point mutation in the upstream homolog arm. Thereafter, the PBacDsRed is excised out, leaving behind the point mutation at the endogenous locus. Point mutation was further confirmed by isolating genomic DNA from a single fly and performing Sanger sequencing.

### Fly husbandry

All the fly stocks and crosses were maintained at 25°C. Egg collection were done in 6 hours’ window and the larvae were reared at 25°C (for mutants) or 29°C (for RNAi knockdown/overexpression/rescue experiments) until dissection. Details of the genotypes analysed in each figure are given in the supplementary data.

### RNA interference experiments

Virgin females of *UAS-dcr2; inscGAL4-UASmCD8-GFP* were crossed to the flies of the following genotypes and progeny of the required genotypes were separated and dissected: *Canton-S, UAS-brm-RNAi, UAS-HA-brm-K804R, UAS-Su(H)-RNAi, UAS-Notch-RNAi, UAS-mam-RNAi, UAS-Dl-RNAi, UAS-Nej-RNAi, UAS-HA-brm, UAS-Moira-RNAi, UAS-Snr1-RNAi, UAS-Bap111-RNAi, UAS-Bap55-RNAi, UAS-Osa-RNAi, UAS-SayP-RNAi, UAS-mam-RNAi, UAS-Polybromo-RNAi, UAS-Bap170-RNAi*.

Six-hour egg collections were done, and the embryos were reared at 25°C for 24 hours before shifting to 29°C till late L3 stage dissection, when the larvae were sexed, separated, and subsequently dissected. The effect of these knockdowns on *F3B3-lacZ* was checked in A8-A10 NBs and was quantified and compared to lacZ levels in A8-A10 NBs that have been blocked for apoptosis by expression of p35 (which served as a control). For *UAS-p35* control experiments, virgin females of *UAS-dcr2; inscGAL4 UASmCD8-GFP; tub-GAL80^ts^*were crossed to the *UAS-p35* flies, and larvae were reared at 18°C for 48-54 hours and then shifted to 29°C until dissection at the LL3 stage. The schematic of the temperature-shift (TS) protocol for these experiments is summarised in Fig. S4C.

### Immunostaining staining image acquisition

Larvae of Late L3 stage of desirable sex were dissected and fixed in 4% paraformaldehyde in 1X PBS and 0.1% TritonX-100 for 30 minutes at room temperature. The following primary antibodies were used: rabbit anti-*Dpn*, 1:2000; rat anti-*Dpn*, 1:1000; rabbit anti-*Grh*, 1:2000, mouse Abd-B (1:500, 1A2E9, DSHB), mouse anti-*NICD* (1:50, C17.9C6, DHSB), mouse anti-*Dl* (1:10, C594.9B, DHSB), mouse anti-*Ct* (1:500, 2B10, DHSB), chicken anti-GFP (1:2000, ab13970, Abcam), chicken anti-β-gal (1:2000, ab9361, Abcam), chicken anti-HA (1:500, ab9111, Abcam), mouse anti-*mam* (1:1000 PMID: 8725234), rat anti-*nej* (1:1000 PMID: 19075001), mouse anti-*Su(H)* (1:500, SC-398453, Santa-Cruz), Rabbit anti-*H3K27ac* (1:1000, ab4729, Abcam). Secondary antibodies conjugated to Alexa fluorophores from Molecular Probes were used: AlexaFluor-405 (1:250); AlexaFluor-488 (1:500); AlexaFluor-555 (1:1000); and AlexaFluor-647 (1:250). The samples were mounted in 70% glycerol and images were acquired with Zeiss LSM 700 and LSM 900 confocal microscopes and processed using ImageJ and Adobe Illustrator CS3. Scale bars are shown in figure legends. Microsoft Excel and GraphPad Prism was used for all the data analysis (unpaired student t-test was done to check significance of the data). At least three technical replicates were done for all the experiments. A minimum of seven VNCs were analyzed across these replicates, and a representative confocal image is shown for each experiment. Since making a z-project for all confocal slices often compromises the signal-to-noise ratio and obscures information, single confocal z-slices are used as representative images across all experiments, and the data are quantified in the graph.

### RNA isolation, reverse transcription and qRT-PCR

RNA extraction was done using 50-60 L3 larval brains of suitable genotype. TRIzol (Thermo Fisher #15596018) was used, followed by chloroform extraction and isopropanol precipitation. Thereafter RNA was resuspended in RNase free water.

For reverse transcription, RNA was treated with DNase (Invitrogen TURBO DNA-free kit AM1907) and then converted to cDNA using Takara cDNA synthesis kit (PrimeScript cDNA Synthesis Kit #6110A). cDNA was diluted five-fold before using for qRT-PCR. cDNA was then subjected to qRT-PCR using Takara SYBR Green (TB Green Premix Ex Taq, ROX Plus #RR42LR) in Himedia LA1074-1NO Insta Q96 - 6.0. Transcript levels were quantified using 2^-ΔΔCt^ method [53] [54], and transcript levels were normalized using GAPDH. Primer sequences are provided in the supplementary data.

### Dam-ID and qRT-PCR

We used NB-specific *inscGAL4* to drive the expression of very low levels of Dam-Brm fusion protein and Dam protein alone. The Brm binding in the genome was assayed by methylation of nearby GATC sequences and normalised over baseline GATC methylation carried out by Dam alone, using RT-PCR. Both controls and test experiments were done three time and average value was taken for comparison.

To check *Brm* binding on 1kb apoptotic enhancer, *UAS-LT3-Dam* and *UAS-LT3-DamBrm* flies [44] were crossed to *UAS-dcr2; inscGAL4 UASmCD8-GFP; tub-GAL80^ts^* virgin female flies. Six hour egg collection was done in 25°C embryos were shifted and reared at 18°C until mL2 stage and then shifted to 29°C (TS as shown in the Fig. S4B). 50-60 brains with VNC from mL3 stage were dissected and collected in 1X PBS and stored in −80°C until used. The tissues were processed as per the published Targeted DamID protocol [55], where genomic DNA was extracted from control and test tissues, digested with Dpn I, followed by adapter ligation and Dpn II digest, thereafter PCR amplification and sonication was done. Subsequently, adaptors were removed, and the sonicated genomic samples were subjected to qRT-PCR. The samples were diluted five-fold before qRT-PCR. Fold enrichment in case of Dam-Brm on *F3B3* was compared to Dam-only using formula 2^Ct^ ^(Dam-only)−Ct^ ^(Dam-*Brm*)^. Sequence of the *F3B3* specific primers used are given in the supplementary data.

### *In vitro* GST-pulldown

The following proteins were used for *in vitro* pulldown: GST-Su(H), 6X His-Brm F1 (1-311aa), His-Brm F2 (304-747aa) and His-Brm F3 (748-1638aa). Bacterial cultures expressing full length GST-tagged *Su(H)* and GST-only were induced for 4 hours with 0.5C:mM IPTG at 18°C and different fragments of His-tagged Brm proteins were induced for 4 hours with 0.25mM IPTG at 37°C. After sonication, lysates were treated with Micrococcal Nuclease (MN) at 37°C for half an hour. GST-Su(H) and GST-only were incubated with GST beads for 2 h at 4°C. Overnight IP was set with bead-bound GST-Su(H) and GST-only and His-tagged Brm F1, F2 and F3 lysates. Bead-bound proteins were separated by denaturing SDS-PAGE and then transferred onto PVDF membranes. The membrane was blocked in 5% milk in Tris-buffered saline with 0.1% Tween 20 (TBST). Primary antibody-mouse anti-His (H1029; Sigma-Aldrich) was diluted 1 in 5000 in 5% milk in TBST, and the blot was incubated overnight at 4°C. HRP conjugated secondary antibody (Peroxidase-AffiniPure Rabbit Anti-Mouse IgG + IgM(HC:+C:L) (1/10,000, 315-035-048; Jackson Immunological Research Laboratory, USA) was used. Visualization was carried out by Enhanced Chemiluminescent femtoLUCENT PLUS HRP chemiluminescent detection (G Biosciences #786003) in Intelligent ImageQuant LAS 500. Representative blots from at least 3 repetitions of the experiments are shown in the figures.

### *In vivo* pulldown assays with whole larvae

50 larvae were homgenized and lysed in RIPA buffer (150mM NaCl, 10mM Tris-HCl pH 7.5, 1mM EDTA, 0.1% Triton X-100, 0.1% SDS, 0.1% Sodium deoxycholate, 1mM PMSF and Protease Inhibitor). Protein A beads were incubated with Rabbit Anti-IgG (Merck 12-370, Lot #2378704) and Rabbit Anti-Brm [40] for 5 hours at 4°C. Pre-cleaned lysates were then set for overnight IP with protein A beads bound to Rabbit Anti-IgG and Rabbit Anti-Brm. Next day beads were washed and a denaturing SDS-PAGE was run. Proteins were then transferred to PVDF membrane and membrane was blocked in 5% milk in Tris-buffered saline with 0.1% Tween 20 (TBST). Primary antibody, mouse anti-mam (1:1000;[56]), rat anti-*nej* (1:1000; [57]), and rat anti-Su(H) (1:2000, MABE982, Merck) were diluted in 5% milk in TBST and incubated overnight at 4°C. HRP conjugated secondary antibody Peroxidase-AffiniPure Rabbit Anti-Mouse IgG + IgM(HC:+C:L) (1:10,000, 315-035-048; Jackson Immunological Research Laboratory, USA), Peroxidase AffiniPure Goat Anti-Rat IgG (H+L) (1:10,000, 112-035-003; Jackson Immunological Research Laboratory, USA) were used. Visualization was carried out by Enhanced Chemiluminescent femtoLUCENT PLUS HRP chemiluminescent detection (G Biosciences #786003) in Intelligent ImageQuant LAS 500. Representative blots from at least three repetitions of the experiments are shown in the figures.

## Supporting information

Highlights

genotypes used

## Acknowledgements

We thank J. Knoblich, A. Brand for reagents; BDSC, VDRC, NIG-Fly, and DGRC - Japan stock centres for fly lines; CDFD animal facility, Bioklone Biotech Pvt. Ltd., Chennai and the Developmental Studies Hybridoma Bank at The University of Iowa for antibodies and TFF at NCBS-CCAMP, Bangalore for transgenic flies. We thank Raviranjan Kumar for making initial observations with Brahma knockdown lines, Prakeerthi Abburi for help with generation of Pygenome tracks for Targeted DamID data (published by Marshall and Brand, 2017) showing Brm binding on different genes for Figure S6 and help with GLM statistical analysis, other members of LNCB for their comments and C. S. Singh for his assistance in various phases of the project. This study was funded by; Department of Science and Technology, India (CRG/2021/003275); Department of Biotechnology, India (BT/PR27455/BRB/10/1647/2018, BT/PR41306/MED/122/ 259/2020 and BT/PR45460/MED/12/952/2022); CDFD core funds; and DST Inspire, India (award to PB) [Reference No IF170651].

## Author contribution

RJ conceptualised the study; PB and VP did the experiments; RJ and PB analysed the data; RJ and PB wrote the manuscript.

During the preparation of this work, the author(s) used Grammarly software to check for spelling, grammar, punctuation, clarity, engagement, and delivery mistakes in English text. After using this tool/service, the corresponding author reviewed and edited the content as needed and takes full responsibility for the content of the publication.

## Data Availability Statement

All relevant data are within the manuscript and its Supporting Information files.

---

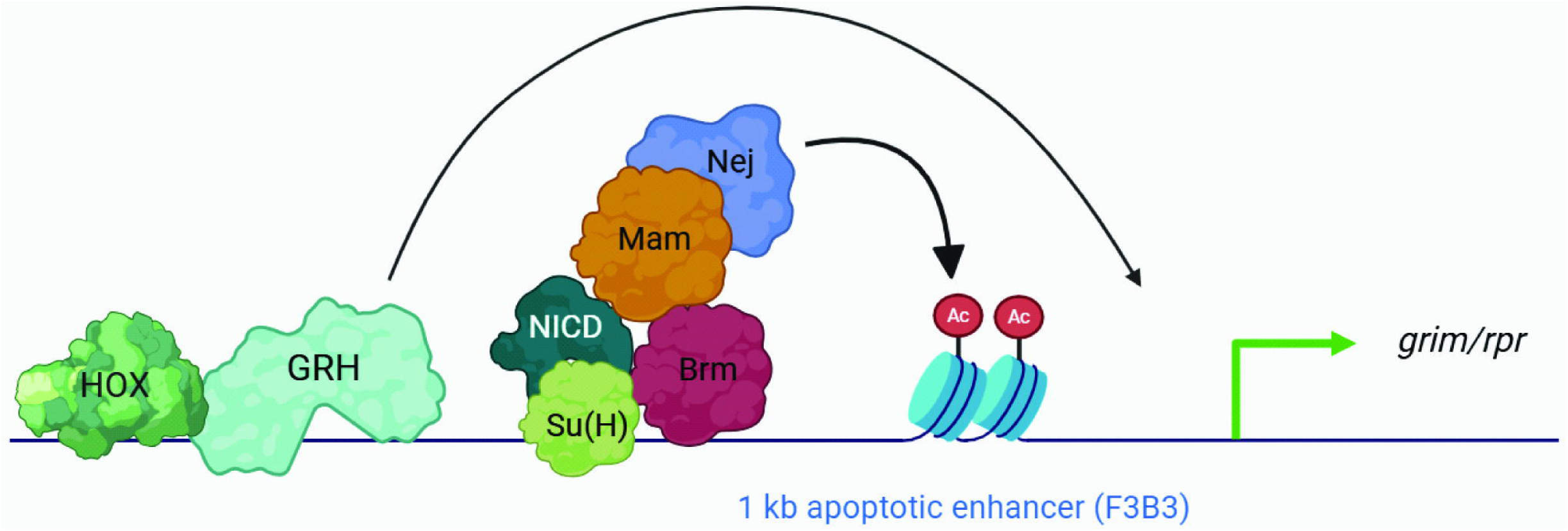

