## SUPPLEMENTARY MATERIAL for "SWI/SNF ATPase Brahma and Notch signaling collaborate with CBP/p300 to regulate neural stem cell apoptosis in *Drosophila* larval central nervous system"

**Supplementary Information**

| **Targeted Dam-ID primers:** | |
| --- | --- |
| F3B3_F1_fwd: CTGTATGTTTGCACGGAGTGGA | |
| F3B3_F1_rev: TTTAAGTTTCGAATAAAGGAAAGATAATG | |
| F3B3_F2_fwd: CCTTTATTCGAAACTTAAAATTTTCTG | |
| F3B3_F2_rev: GAAAATTGTGTTTTTGTAAAAGGATTAG | |
| F3B3_F3_fwd: CAAAAACACAATTTTCTAAGATCCACTG | |
| F3B3_F3_rev: ATATATCCGCACATGCAACCCTC | |
| F3B3_F4_fwd: GGGTTGCATGTGCGGATATATATA | |
| F3B3_F4_rev: TGACGCGCTATGCGACAATTTGAG | |
| F3B3_F5_fwd: AAATTGTCGCATAGCGCGTCAAAT | |
| F3B3_F5_rev: CCCTTTCGTTTGGTCCTTTGTTTG | |
| mβ tr_forward AGAAGTGAGCAGCAGCCATC | |
| mβ tr_reverse GCTGGACTTGAAACCGCACC | |
| mβ enh_forward AGAGGTCTGTGCGACTTGG | |
| mβ enh_reverse GGATGGAAGGCATGTGCT | |
| mβ‐mα igr_forward AAGCCAGTGGACTCTGCTCT | |
| mβ‐mα igr_reverse TGATCTCCAAGCGGAGTATG | |
| **RT-qPCR Experiments:** | |
| Grim_fwd TTAGTTCTCCTTGGAGGTGG | |
| Grim_rev TTCCGTGCCGCTGGGAAG | |
| Rpr_fwd ATGGCAGTGGCATTCTACATAC | |
| Rpr_rev TCATTGCGATGGCTTGCGATATTTG | |
| Delta_fwd CGTGAAAAGGAGAAGACTCCG | |
| Delta_rev CGCACATGGTTTTGCAATTCGC | |
| Brm_fwd CACGGAATGCCACCCAATG | |
| Brm_rev GAGTAGCGAGGATCCTCTTG | |
| GAPDH_fwd GGTGATCAACGACAACTTCG | |
| GAPDH_rev CCAGTGGAAGCTGGAATGAT | |
| **BAP and PBAP RNAi lines** | **Fly Line number** |
| UAS-Moira-RNAi(1) | VDRC 6969 |
| UAS-Moira-RNAi(2) | VDRC 110712 |
| UAS-Snr-RNAi(1) | BDSC 32372 |
| UAS-Snr-RNAi(2) | VDRC 108599 |
| UAS-Bap111-RNAi(1) | VDRC 37682 |
| UAS-Bap111-RNAi(2) | VDRC 104361 |
| UAS-Bap55-RNAi | VDRC 24703 |
| UAS-Osa-RNAi(1) | BDSC 31266 |
| UAS-Osa-RNAi(2) | VDRC 7810 |
| UAS-Polybromo-RNAi(1) | BDSC 32840 |
| UAS-Polybromo-RNAi(2) | VDRC 108618 |
| UAS-Polybromo-RNAi(3) | VDRC 330189 |
| UAS-SayP-RNAi | VDRC 105946 |
| UAS-Bap170-RNAi(1) | BDSC 26308 |
| UAS-Bap170-RNAi(2) | VDRC 34582 |

**List of genotypes analyzed in main and supplementary figures**

**Main Figures:**

Fig-1B-B”: *UAS-dcr2/+; inscGAL4-UASmCD8GFP/+; +/+*

Fig-1C-C”: *UAS-dcr2/+; inscGAL4-UASmCD8GFP/+; UAS-brm-RNAi/+*

Fig-1D-D”: *UAS-dcr2/+; inscGAL4-UASmCD8GFP/+; +/+*

Fig-1E-E”: *UAS-dcr2/+; inscGAL4-UASmCD8GFP/UAS-HA-brm-K804R; +/+*

Fig-1G-G”: *Canton-S*

Fig-1H-H’’: *w*/*w*; *+/+; +/brm^[d00415]^*

Fig-1I-I’’: *w*/*w*; *+/+; +/brm^[I21]^*

Fig-1J-J’’: *w*/*w; +/+; brm^[d00415]^*/*brm^[I21]^*

Fig-1K-K’’: *w*/*w; +/+; +/brm^K804R^*

Fig-1L-L’’: *w*/*w; +/+; brm^[d00415]^/brm^K804R^*

Fig-2A-A’’*: UAS-dcr2/+; inscGAL4-UASmCD8GFP/+; UAS-p35- tub-GAL80^ts^/+*

Fig-2B-B’’: *UAS-dcr2/+; inscGAL4-UASmCD8GFP/+; UAS-brm-RNAi/+*

Fig-2D-D’’: *UAS-dcr2/+; inscGAL4-UASmCD8GFP/+; UAS-p35- tub-GAL80^ts^/+*

Fig-2E-E’’: *UAS-dcr2/+; inscGAL4-UASmCD8GFP/+; UAS-brm-RNAi/+*

Fig-2G-G’’ *w*/*w*; *mγ-GFP/+; worniuGAL4-UAS-dcr2/UAS-p35- tub-GAL80^ts^*

Fig-2H-H’’ *w*/*w*; *mγ-GFP /+; worniuGAL4-UAS-dcr2/UAS-brm-RNAi*

Fig-2I-I’’: *UAS-dcr2/+; inscGAL4-UASmCD8GFP/+; UAS-p35- tub-GAL80^ts^/+*

Fig-2J-J’’: *UAS-dcr2/+; inscGAL4-UASmCD8GFP/+; UAS-brm-RNAi/+*

Fig-2K-K’’: *UAS-dcr2/+; inscGAL4-UASmCD8GFP/+; UAS-p35- tub-GAL80^ts^/+*

Fig-2L-L’’: *UAS-dcr2/+; inscGAL4-UASmCD8GFP/+; UAS-brm-RNAi/+*

Fig-3A-A’’: *Canton-S*

Fig-3B-B’’: *w*/*w*; *+/+; +/brm^2^*

Fig-3C-C’’: *N^[55e11]^*/*w*; *+/+; +/+*

Fig-3D-D’’: *N^[55e11]^*/*w*; *+/+; +/brm^2^*

Fig-3F: *UAS-dcr2/+; inscGAL4-UASmCD8GFP/+; +/+*

*UAS-dcr2/+; inscGAL4-UASmCD8GFP/+; UAS-brm-RNAi/+*

*UAS-dcr2/+; inscGAL4-UASmCD8GFP/UAS-Su(H)-RNAi; +/+*

*UAS-dcr2/+; inscGAL4-UASmCD8GFP/UAS-Su(H)-RNAi; UAS-brm-RNAi/+*

Fig-3I: w/*UAS-dcr2/+; inscGAL4-UASmCD8GFP/+; UAS-LT3-Dam/ tub-GAL80^ts^*

w/*UAS-dcr2/+; inscGAL4-UASmCD8GFP/+; UAS-mcherry-Dam-Brm/ tub-GAL80^ts^*

Fig-4A: *w*/*w*; *F3B3-LacZ/+; worniuGAL4-UAS-dcr2/UAS-p35-tub-GAL80^ts^*

*w*/*w*; *F3B3-LacZ/+; worniuGAL4-UAS-dcr2/UAS-brm-RNAi*

*w*/*w*; *F3B3-LacZ/ UAS-brm-K804R; worniuGAL4-UAS-dcr2/+*

*w*/*w*; *F3B3-LacZ/+; worniuGAL4-UAS-dcr2/UAS-Notch-RNAi*

*w*/*w*; *F3B3-LacZ/+; worniuGAL4-UAS-dcr2/UAS-Mam-RNAi*

Fig-4B: Canton-S

*w/w; +/+; Δ717/ Δ717*

Fig-4C, D, E:	*UAS-dcr2/+; inscGAL4-UASmCD8GFP/+; +/+*

*UAS-dcr2/+; inscGAL4-UASmCD8GFP/UAS-Delta-RNAi; +/+*

*UAS-dcr2/+; inscGAL4-UASmCD8GFP/+; UAS-brm-RNAi/+*

*UAS-dcr2/+; inscGAL4-UASmCD8GFP/ UAS-Delta-RNAi; UAS-brm-RNAi/+*

Fig-4F: *UAS-dcr2/+; inscGAL4-UASmCD8GFP/+; +/+*

*w/w; UAS-HA-brm/+; +/+*

*UAS-dcr2/+; inscGAL4-UASmCD8GFP/+; UAS-Notch-RNAi/+*

*UAS-dcr2/+; inscGAL4-UASmCD8GFP/UAS-HA-brm; UAS-Notch-RNAi/+*

*UAS-dcr2/+; inscGAL4-UASmCD8GFP/+; UAS-Su(H)-RNAi/+*

*UAS-dcr2/+; inscGAL4-UASmCD8GFP/UAS-HA-brm; UAS-Su(H)-RNAi/+*

*UAS-dcr2/+; inscGAL4-UASmCD8GFP/+; UAS-Mam-RNAi/+*

*UAS-dcr2/+; inscGAL4-UASmCD8GFP/UAS-HA-brm; UAS-Mam-RNAi/+*

Fig-5A-A’’: *UAS-dcr2/+; inscGAL4-UASmCD8GFP/+; +/+*

Fig-5B-B’’: *UAS-dcr2/+; inscGAL4-UASmCD8GFP/+; UAS-nej-RNAi/+*

Fig-5E-E’’: *UAS-dcr2/+; inscGAL4-UASmCD8GFP/+; UAS-p35tub-GAL80^ts^/+*

Fig-5F-F’’: *UAS-dcr2/+; inscGAL4-UASmCD8GFP/+; UAS-nej-RNAi/+*

Fig-5H-H’’: *UAS-dcr2/+; inscGAL4-UASmCD8GFP/+; UAS-p35tub-GAL80^ts^/+*

Fig-5I-I’’: *UAS-dcr2/+; inscGAL4-UASmCD8GFP/+; UAS-brm-RNAi/+*

Fig-5C-C’’: *UAS-dcr2/+; inscGAL4-UASmCD8GFP/+; UAS-p35tub-GAL80^ts^/+*

Fig-5D-D’’: *UAS-dcr2/+; inscGAL4-UASmCD8GFP/+; UAS-nej-RNAi/+*

Fig-5J-J’’: *UAS-dcr2/+; inscGAL4-UASmCD8GFP/+; UAS-brm-RNAi/+*

Fig-5K-K’’: *UAS-dcr2/+; inscGAL4-UASmCD8GFP/UAS-HA-brm; UAS-nej-RNAi/+*

Fig-5G: *w*/*w*; *F3B3-LacZ/+; worniuGAL4-UAS-dcr2/UAS-p35tub-GAL80^ts^*

*w*/*w*; *F3B3-LacZ/+; worniuGAL4-UAS-dcr2/UAS-nej-RNAi*

**Supplementary Figures:**

Fig-S1B-B’’: *UAS-dcr2/+; inscGAL4-UASmCD8GFP/+; +/+*

Fig-S1C-C’’: *UAS-dcr2/+; inscGAL4-UASmCD8GFP/+; UAS-Moira-RNAi/+*

Fig-S1D-D’’: *UAS-dcr2/+; inscGAL4-UASmCD8GFP/UAS-Moira-RNAi; +/+*

Fig-S1E-E’’: *UAS-dcr2/+; inscGAL4-UASmCD8GFP/+; UAS-Snr1-RNAi/+*

Fig-S1F-F’’: *UAS-dcr2/+; inscGAL4-UASmCD8GFP/UAS-Snr1-RNAi; +/+*

Fig-S1G-G’’: *UAS-dcr2/+; inscGAL4-UASmCD8GFP/+; UAS-Bap111-RNAi/+*

Fig-S1H-H’’: *UAS-dcr2/+; inscGAL4-UASmCD8GFP/UAS-Bap111-RNAi; +/+*

Fig-S1I-I’’: *UAS-dcr2/+; inscGAL4-UASmCD8GFP/UAS-Bap55-RNAi; +/+*

Fig-S1J-J’’: *UAS-dcr2/+; inscGAL4-UASmCD8GFP/+; UAS-Osa-RNAi/+*

Fig-S1K-K’’: *UAS-dcr2/+; inscGAL4-UASmCD8GFP/+; UAS-Osa-RNAi/+*

Fig-S1L-L’’: *UAS-dcr2/+; inscGAL4-UASmCD8GFP/+; UAS-Polybromo-RNAi/+*

Fig-S1M-M’’: *UAS-dcr2/+; inscGAL4-UASmCD8GFP/UAS-Polybromo-RNAi; +/+*

Fig-S1N-N’’: *UAS-dcr2/+; inscGAL4-UASmCD8GFP/UAS-Polybromo-RNAi; +/+*

Fig-S1O-O’’: *UAS-dcr2/+; inscGAL4-UASmCD8GFP/UAS-SayP-RNAi; +/+*

Fig-S1P-P’’: *UAS-dcr2/+; inscGAL4-UASmCD8GFP/+; UAS-Bap170-RNAi/+*

Fig-S1Q-Q’’: *UAS-dcr2/+; inscGAL4-UASmCD8GFP/+; UAS-Bap170-RNAi/+*

Fig-S2A-A’’: *UAS-dcr2/+; inscGAL4-UASmCD8GFP/+; +/+*

Fig-S2B-B’’: *UAS-dcr2/+; inscGAL4-UASmCD8GFP/+; UAS-Notch-RNAi/+*

Fig-S2C-C’’: *UAS-dcr2/+; inscGAL4-UASmCD8GFP/UAS-Su(H)-RNAi; +/+*

Fig-S2D-D’’: *UAS-dcr2/+; inscGAL4-UASmCD8GFP/+; UAS-Mam-RNAi/+*

Fig-S2F-F’’: *w*/*w*; *mγ-GFP/+; worniuGAL4-UAS-dcr2/UAS-p35 tub-GAL80^ts^*

Fig-S2G-G’’ *w*/*w*; *mγ-GFP/UAS-Su(H)-RNAi; worniuGAL4-UAS-dcr2/+*

Fig-S2H-H’’ *w*/*w*; *mγ-GFP/+; worniuGAL4-UAS-dcr2/UAS-Mam-RNAi*

Fig-S5A-A’’: *UAS-dcr2/+; inscGAL4-UASmCD8GFP/+; UAS-p35 tub-GAL80^ts^/+*

Fig-S5B-B’’: *UAS-dcr2/+; inscGAL4-UASmCD8GFP/UAS-Su(H)-RNAi; +/+*

Fig-S5C-C’’: *UAS-dcr2/+; inscGAL4-UASmCD8GFP/+; UAS-brm-RNAi/+*

Fig-S5D-D’’: *UAS-dcr2/+; inscGAL4-UASmCD8GFP/UAS-Su(H)-RNAi; UAS-brm-RNAi/+*
