## Supplementary material for "SWI/SNF ATPase Brahma and Notch signaling collaborate with CBP/p300 to regulate neural stem cell apoptosis in *Drosophila* larval central nervous system": Highlights

Brahma regulates Drosophila NSC apoptosis by a two-tier regulation.

Brahma regulates expression of Abd-B, Grainyhead, Notch, Mastermind and CBP/p300 (Nejire).

Brahma and Su(H) physically collaborate on apoptotic enhancer to regulate the *RHG* family of apoptotic genes.

This collaboration recruits CBP/p300 to acetylate nucleosome at H3K27, thereby opening the chromatin to facilitate RHG gene transcription.
