## Supplementary figures and images for "SWI/SNF ATPase Brahma and Notch signaling collaborate with CBP/p300 to regulate neural stem cell apoptosis in *Drosophila* larval central nervous system"

### FigS1

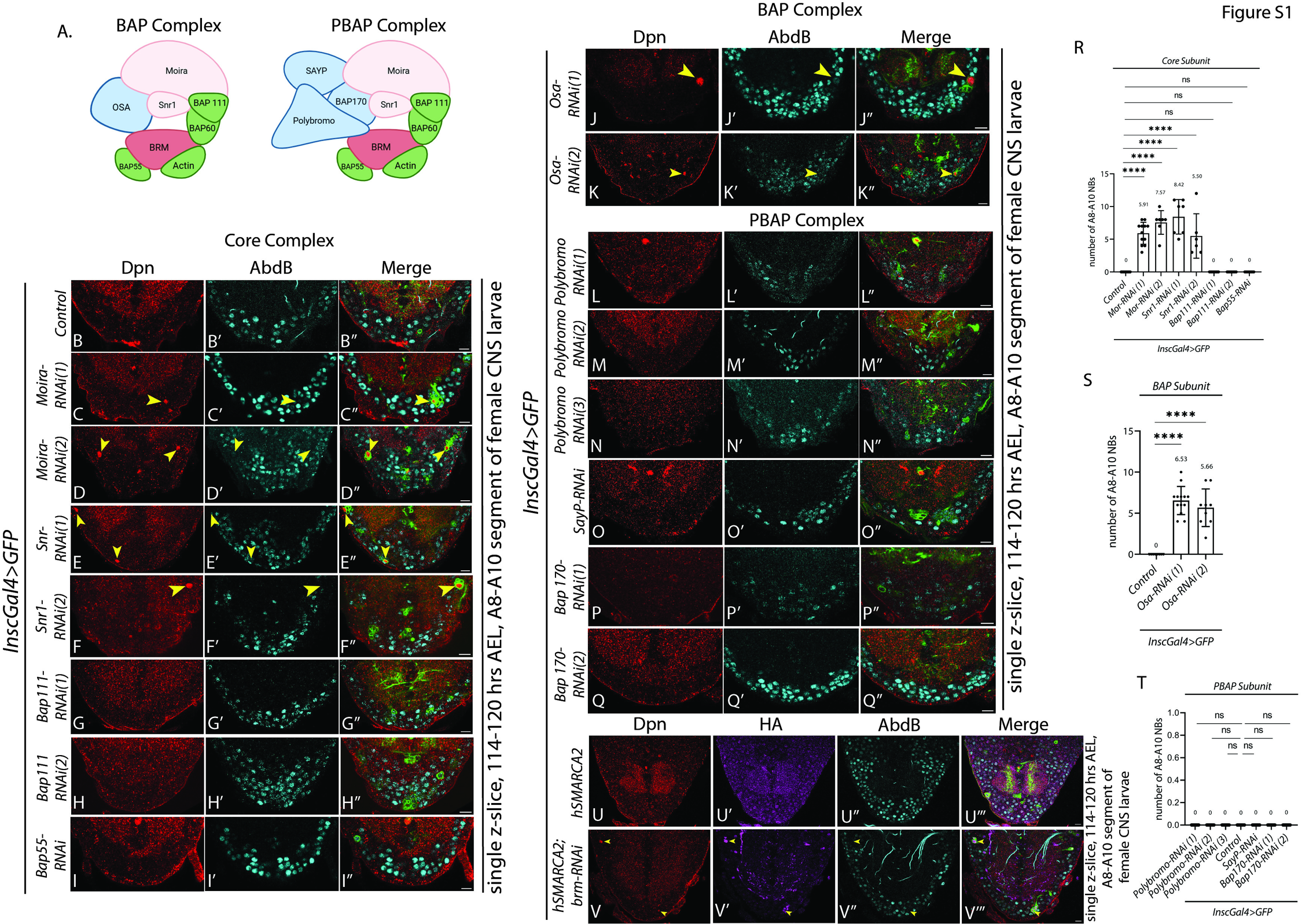

### FigS2

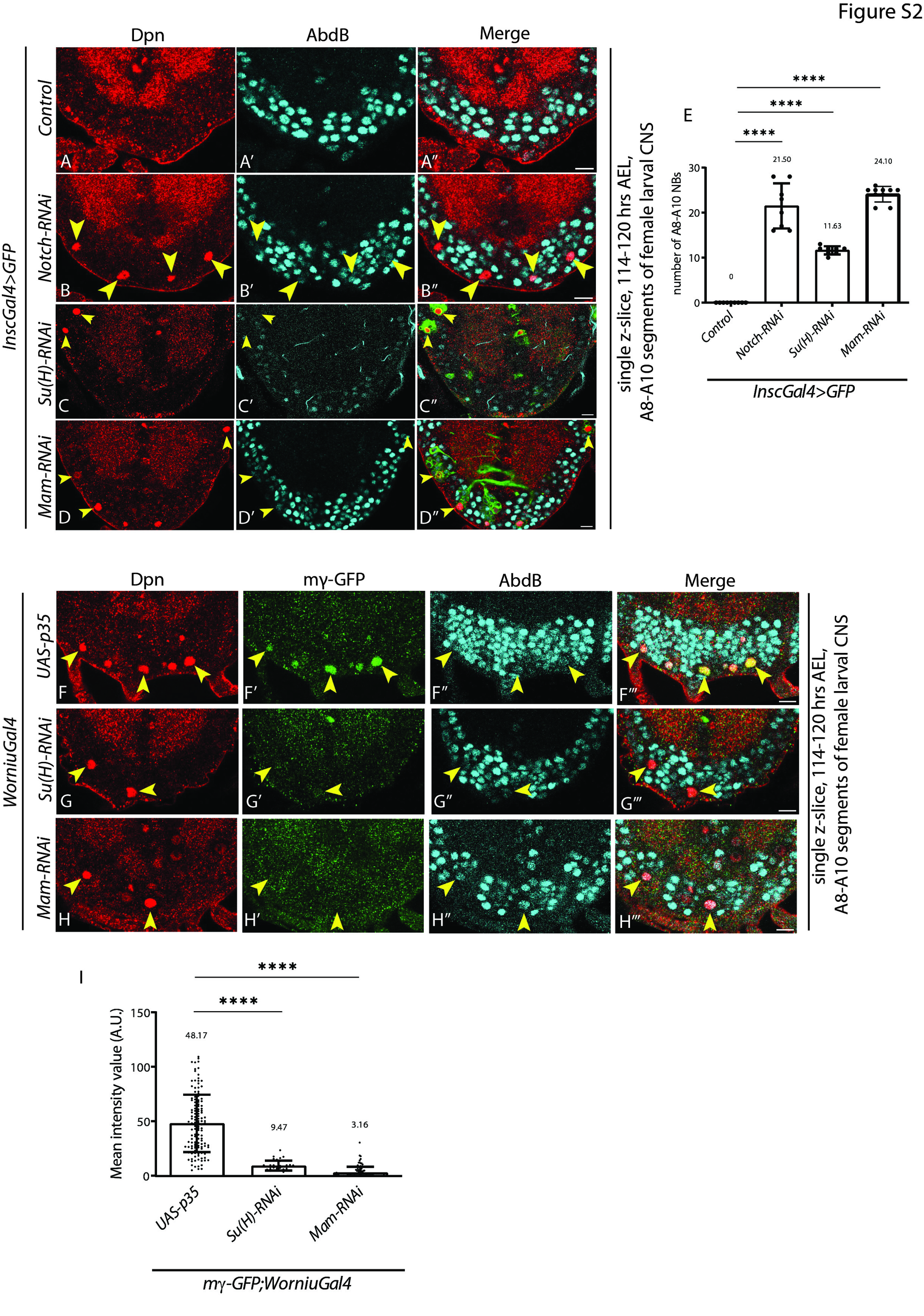

### FigS3

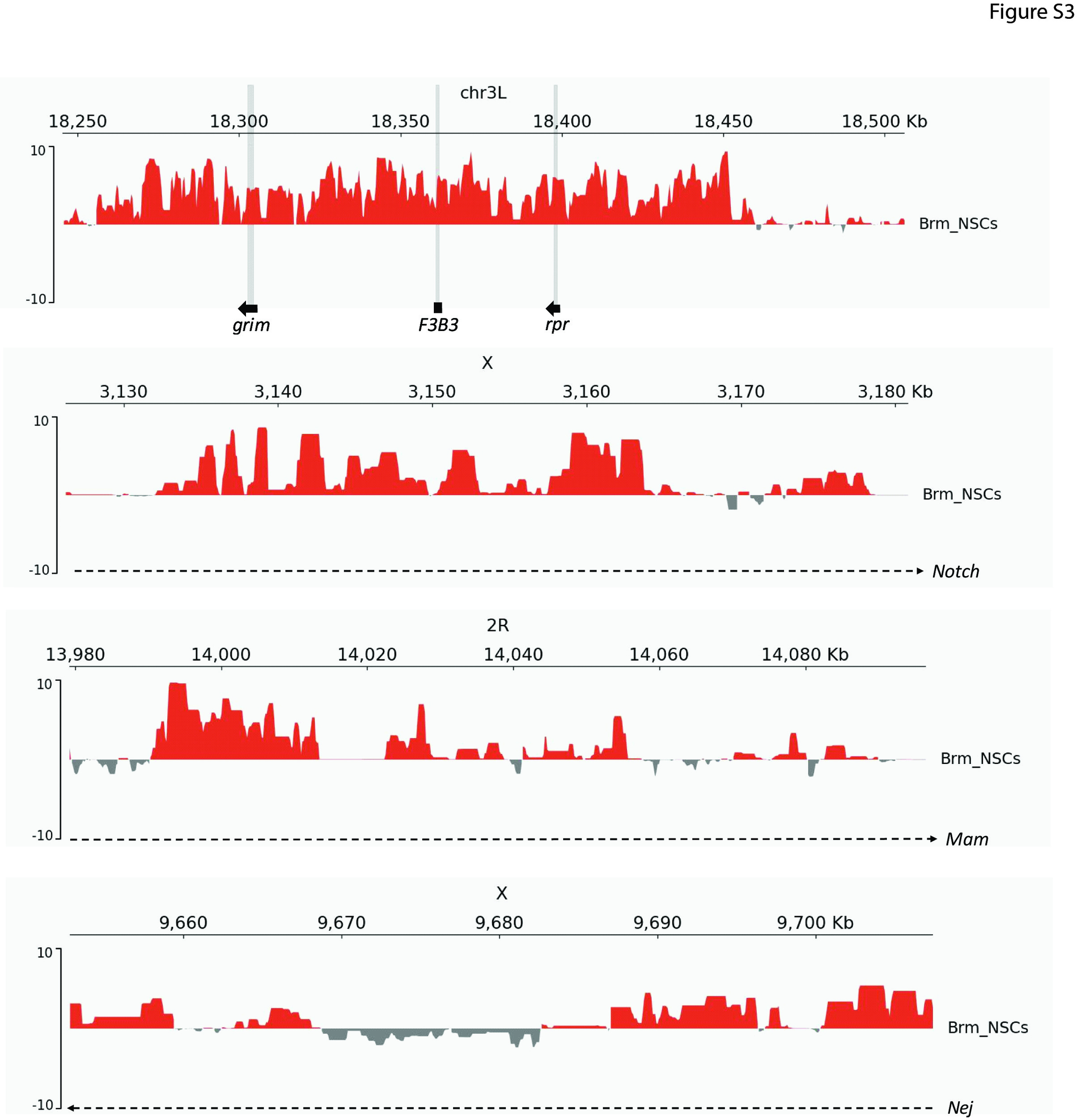

### FigS4

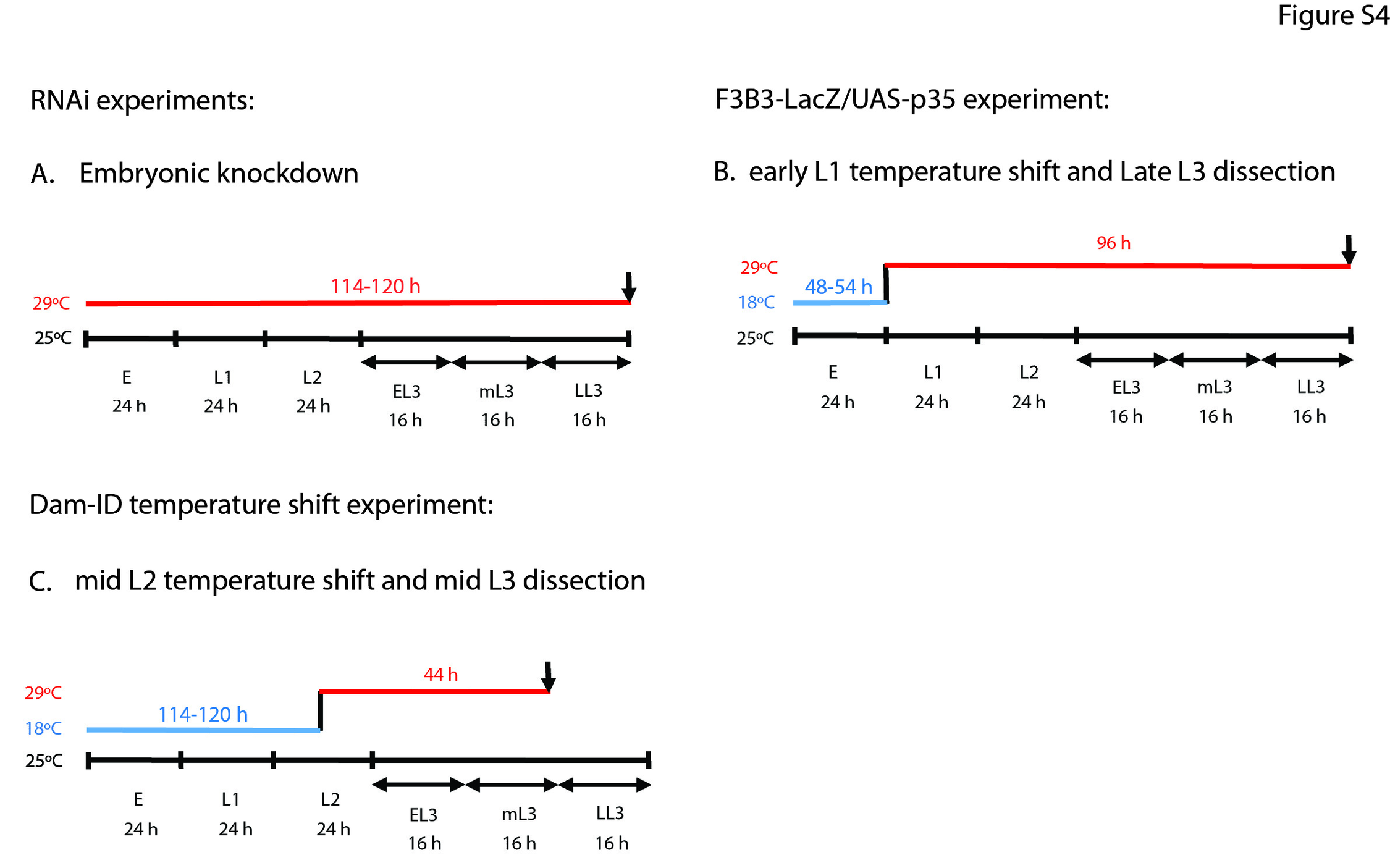

### FigS5

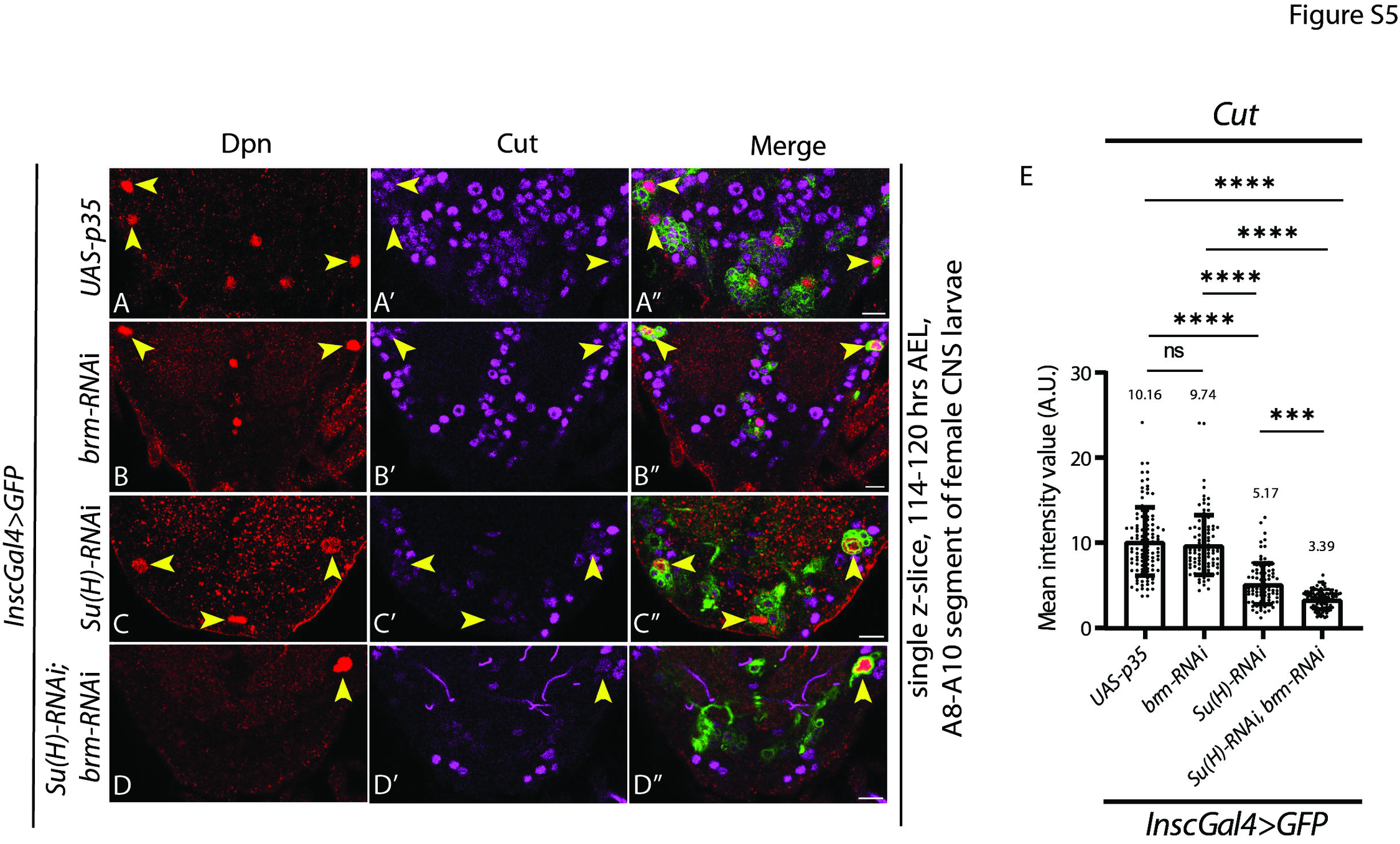

### FigS6

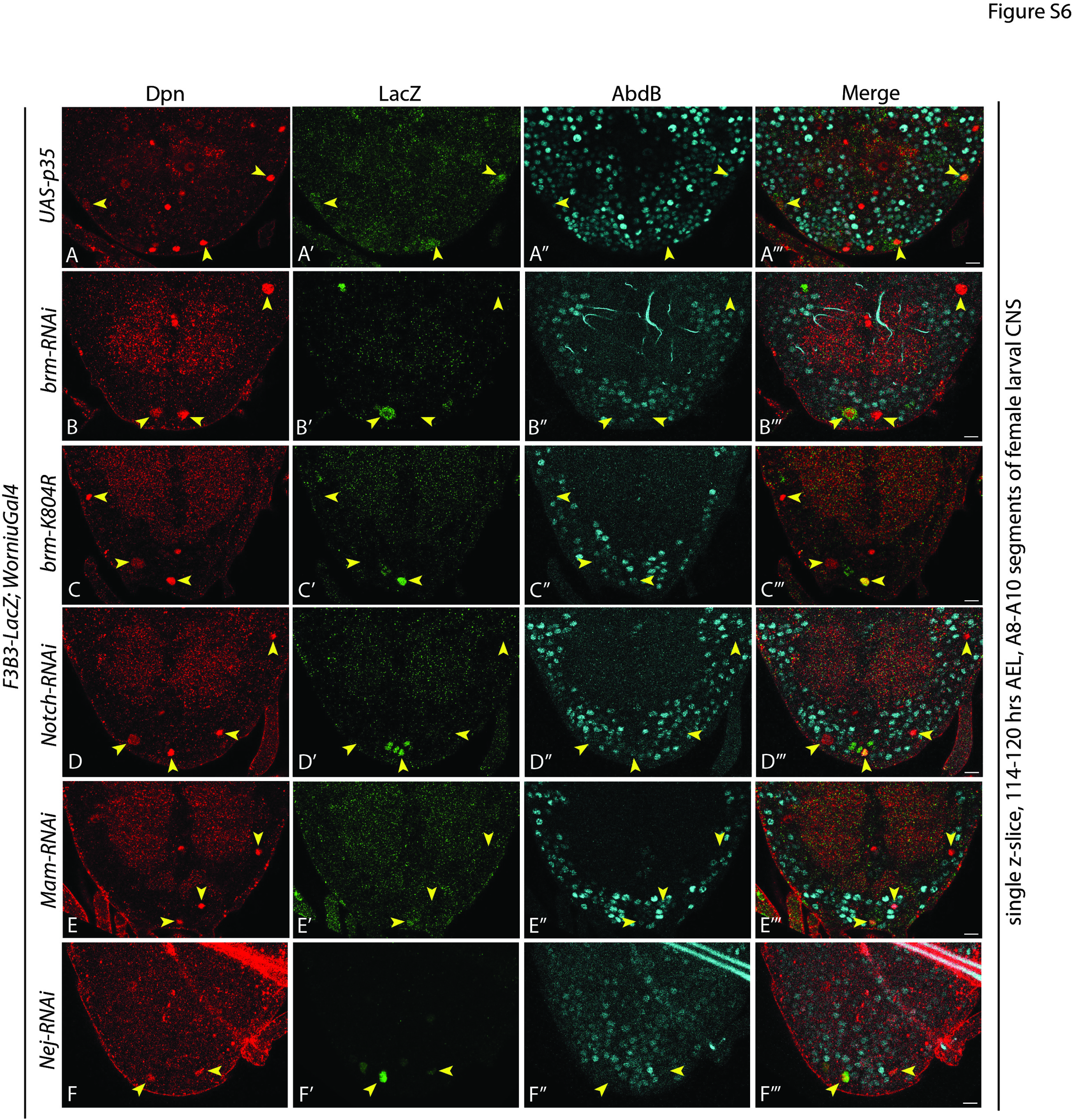

### FigS7

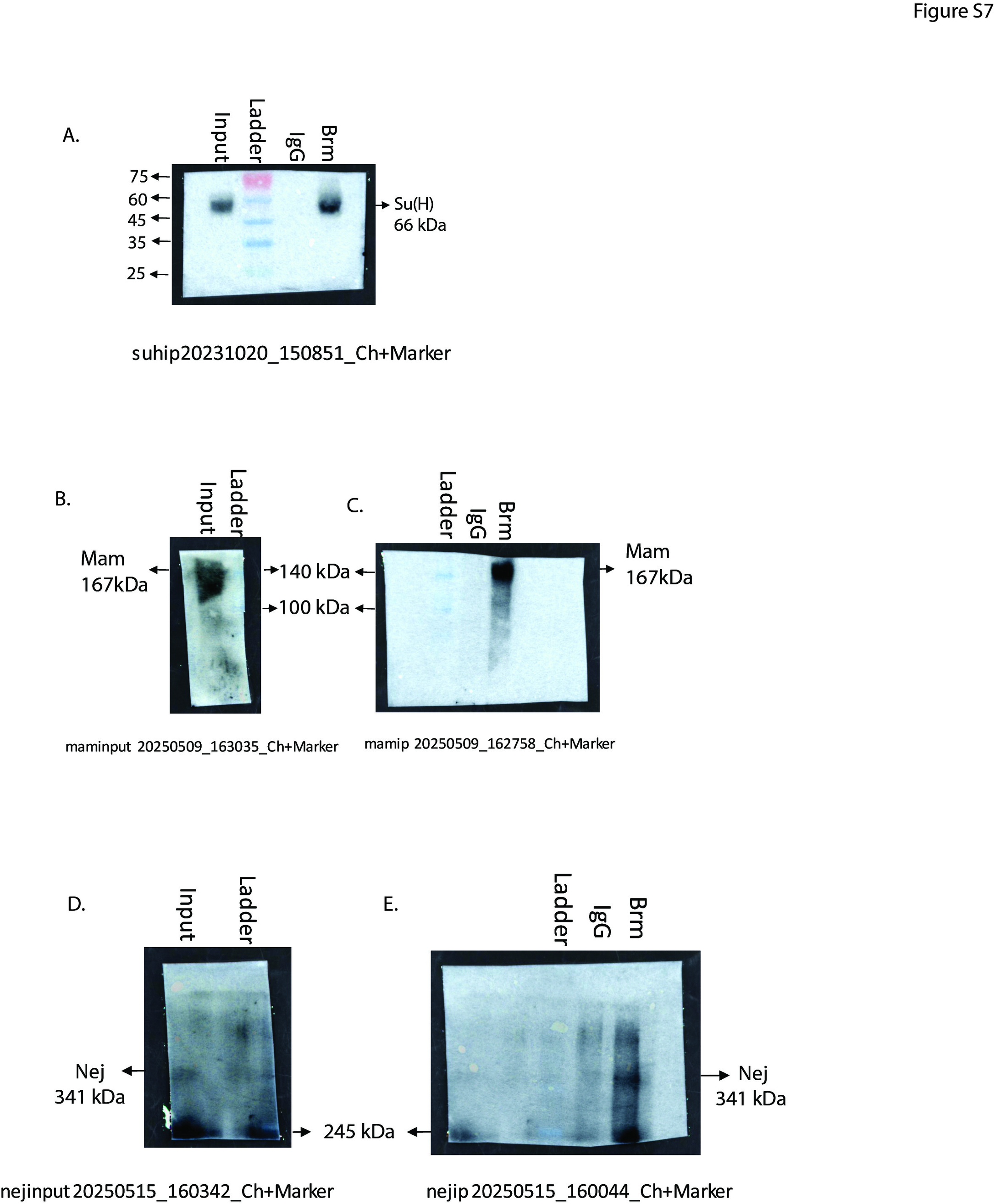
